# High nucleotide similarity of three *Copia* lineage LTR retrotransposons among plant genomes

**DOI:** 10.1101/2022.02.23.481133

**Authors:** Simon Orozco-Arias, Mathilde Dupeyron, David Gutiérrez-Duque, Reinel Tabares-Soto, Romain Guyot

## Abstract

Transposable elements (TEs) are mobile genetic elements found in the majority of eukaryotic genomes. Because of their mobility in the host genome, TEs can deeply impact the structure and evolution of chromosomes and can induce mutations affecting coding genes. In response to these potential threats, host genomes use various processes to repress the TE expression, leading to an arm-race between TEs for their persistence and host genomes for their protection. In plants, the major group of TEs is the Long Terminal Repeats retrotransposons (LTR-RT). They are classified into superfamilies (*Gypsy*, *Copia*) and sub-classified into lineages according to similarities, structures and presence of coding domains. Among the different ways LTR-RTs can proliferate, horizontal transfer (HT), defined as the nonsexual transmission of nuclear and plastid genetic material between species, is a process allowing LTR-RTs to invade a new genome. Although this phenomenon was considered rare in eukaryotic organisms, recent studies demonstrate numerous potential transfers of LTR-RTs, suggesting that HT may be more frequent than initially estimated.

This study aims to determine which LTR-RT lineages are shared with high similarity among 69 reference genomes that represent the major groups of green plants. We first identified and classified 88,450 LTR-RTs and determined 143 cases of high similarities between pairs of genomes. Most of them involved three *Copia* lineages (Oryco/Ivana, Retrofit/Ale and Tork/Tar/Ikeros) and very few of them included the *Gypsy* superfamily. Interestingly, a detailed analysis of three high similarities involving the Tork/Tar/Ikeros group of lineages indicates a patchy distribution of the elements and phylogenetic incongruities, indicating they originated from potential HTs. Overall, our results demonstrate that three specific lineages of *Copia* share outstanding similarity between very distant species and may probably be involved in horizontal transfer mechanisms.

## INTRODUCTION

Transposable elements (TEs) are mobile elements able to move from one locus to another. Their mobility can induce mutations, introduce phenotypic novelties, play key roles in environmental adaptation (Li et al. 2018), and increase their copy number. These elements can also interfere with transcription activities of neighboring genes (Bourque et al. 2018). TEs are classified into two classes according to their replication mechanisms: Class I, called retrotransposons, for TEs moving through an RNA intermediate, (also called “copy and paste” mechanism) and Class II, or transposons, for TEs moving through a DNA intermediate (also called “Cut and Paste” mechanism) (Wicker et al. 2007). Due to the numerous and potentially harmful consequences of their mobility, genomes developed several ways of fighting against the TE proliferation, for example through epigenetic silencing (Slotkin et Martienssen 2007). TEs persistence can be seriously challenged, and a few possibilities remain for them to survive.

One of these possibilities is escaping a genome to invade a new one through horizontal transfer (Blumenstiel 2019). Horizontal transfers (HTs) are defined as the nonsexual transmission of nuclear and plastid genetic material between species. Although this phenomenon was considered uncommon (Panaud 2016), numerous HTs have been demonstrated in the last years in prokaryotes (Diao, Freeling, et Lisch 2005) and eukaryotes (Aubin, El Baidouri, et Panaud 2021). These transfers could contribute to the evolution and adaptation of species due to the diversity of the new acquired genes (Roulin et al. 2009; Acuna et al. 2012; Rice et al. 2013). HTs can involve nuclear and plastid genes (Aubin, El Baidouri, et Panaud 2021), while several studies have also reported the transfer of TEs (or HTT) (Roulin et al. 2007), which sometimes leads to deep genotypic and phenotypic consequences (Gilbert et Feschotte 2018).

In plant genomes, Long Terminal Repeat retrotransposons (LTR-RT) are the most frequent elements (Grandbastien 2015). These can represent up to 80% of the genome size, such as in wheat or barley. LTR-retrotransposons are classified into *Copia* or *Gypsy* superfamilies according to the internal organization of the coding domains (Gao et al. 2012). Each *Copia* and *Gypsy* superfamily is sub-classified into lineages (Wicker et al. 2007) based on specific coding regions and overall structure similarities (Carlos Llorens et al. 2009). Although LTR-RT are frequently compared to retroviruses, the former are endogenous genome elements generally lacking the genetic material to leave the cell and infect other organisms. In plant genomes, *Copia* and *Gypsy* are sub-classified into 16 and 14 lineages, respectively, and some of these are closely related or others restricted to very few species known to date (Neumann et al. 2019). Now, bioinformatic tools allow the precise and rapid classification of LTR-RT based on domain similarities (Orozco-Arias et al. 2018). HTs can be assessed through different methodologies that search for phylogenetic incongruence between TEs and the host genome, the «patchy distribution of conservation» among organisms, and, indeed, the high nucleotide similarity of these elements between distantly related species (Wallau, Ortiz, et Loreto 2012). More than 2,800 HTTs have been documented in eukaryotes, mainly among insect genomes (2,248 HTTs) (Peccoud et al. 2017) as well as between mammals and tetrapods (Pace et al. 2008); fungi and plants (Novikova, Smyshlyaev, et Blinov 2010; Wang et al., 2020); arthropods and conifers (Lin, Faridi, et Casola 2016); bivalves and other aquatic species (Metzger et al. 2018); and birds and nematodes (Suh et al. 2016). A significant number of LTR retrotransposons HTs have been detected in plants (El Baidouri et al. 2014). However, no information has been reported about the classification down to the level of the precise lineage of LTR-RTs involved in these HTs.

To understand the evolution of LTR-RT lineages in plant and to explore the potential of some lineages to be involved in HT, we conducted an *in-silico* reassessment of LTR-RTs shared with high sequence similarity among 69 genomes of green plants, including angiosperm and non-angiosperm species. The analysis of 88,450 LTR-RTs indicates that three *Copia* lineages share high sequence similarity between distantly related species. This finding raises several questions such as, is there more HTs than estimated so far in the plant genomes? What could be the relationship between the success of HT of LTR-RTs in plant genomes and these specific lineages? and what is their capacity to persist in a genome and invade a new one?

## MATERIALS AND METHODS

### Genomic data

We downloaded the genomes of 69 species from 34 plant families (**Supplementary Table 1**), representing a total of 68.4 Gb of data. The relationships between the species used in this analysis are illustrated in CoGePEDIA (https://genomevolution.org/wiki/index.php/Sequenced_plant_genomes, accessed 08/29/2019), Timetree (http://www.timetree.org) and summarized in **Supplementary Figure 1**.

### LTR-RT identification and classification

The genome sequences were first processed with LTR_STRUC (McCarthy et McDonald 2003) for *de novo* prediction of LTR retrotransposons. The 88,450 predicted LTR-RTs were classified using Inpactor ((Orozco-Arias et al. 2018), https://github.com/simonorozcoarias/Inpactor) into superfamilies (i.e. *Copia* or *Gypsy*) and lineages (**Supplementary Table 2**). The lineage names were re-organized according to the GyDB ((C. Llorens et al. 2011), http://gydb.org) and REXDB classifications (Neumann et al. 2019). In some cases, synonymy of Gypsydb and REXDB was used as follow: Del/Tekay, Ivana/Oryco, Ale/Retrofit and closely related lineages defined in REXDB were grouped to correspond to the Gypsydb classification as follow: Tork/Tar/Ikeros.

### Comparison of predicted LTR retrotransposons

Pairwise comparisons of the predicted LTR retrotransposons were performed using the BLASTn algorithm (with-evalue 1e-4 and the other parameters by default). The LTR-RT superfamilies (i.e. *Copia* and *Gypsy*) and lineages from one species were compared to those from the other species. The highest BLASTn “bit-score” for each pairwise comparison was kept and displayed on a graphical heatmap matrix using gnuplot (http://gnuplot.sourceforge.net). This score is directly associated with each pair of residues according to a nucleotide similarity matrix (Match/Mismatch Scores: 2-3; Gap cost: Open 5, Extension 2). In particular, a higher score indicates a higher identity between the aligned residues. The upper limit of similarity on the graphical heatmap was set to BLASTn bit-score values of 3,500, 3,700, 4,000, 5,000 and 6,000 (**Supplementary Figure 2**) to determine the most informative value. The results were organized according to the order shown in the representation of the phylogeny of the plant species (**Supplementary Figure 1)**.

### Characterization of highly similar LTR retrotransposon elements

The shared LTR retrotransposons between two species were carefully analyzed and characterized to confirm the high similarity observed using the BLASTn bit-score matrix. The LTR-RTs were annotated using Artemis (Rutherford et al. 2000) and pairwise compared using dotter (Sonnhammer et Durbin 1995) and Stretcher from EMBOSS (Rutherford et al. 2000). RT phylogenetic trees were obtained based on the results from Step 4 of Inpactor (Orozco-Arias et al. 2018) and edited with Figtree (http://tree.bio.ed.ac.uk/software/figtree/). Full-length LTR-RT sequences were aligned using nucmer with the following parameters: -l 10 -c 10 --nosimplify –maxmatch, and displayed with mummerplot from the MUMer package (Kurtz et al. 2004). For seven plant species, namely poplar, coffee, cannabis, orchid, banana, soybean, and kiwifruit (**Supplementary Table 3)**, nuclear coding sequences (CDS) were retrieved from public databases and pairwise compared using BLASTn (setting the parameter -qcov_hsp_perc equal to 60).

### Nuclear phylogeny of plant species

The nuclear phylogeny of plant species used in this study was established using near-universal single-copy ortholog genes recovered with Busco (BUSCO v5.2.2) for the 69 plant genomes. The phylogeny was carried out using the BUSCO_Phylogenomics pipeline (https://github.com/jamiemcg/BUSCO_phylogenomics; (McGowan et al. 2020)) and the super tree approach. The phylogeny of LTR-RT was carried out using Reverse Transcriptase domains (RT). RT domains were extracted from the predicted and classified LTR-RTs similarly to (Ming et al. 2015). RT classified as TORK (1235 sequences) were aligned with MAFFT V. 7.471, (Katoh et Standley 2013) and FastTree V.2.1.10, (Price, Dehal, et Arkin 2010) was used to perform the phylogenetic analysis.

## RESULTS

### Screening of LTR retrotransposons conservations across plant genomes

LTR retrotransposons (LTR-RTs) were mined from 69 available plant genomes (angiosperm and non-angiosperm species, Viridiplantae; **Supplementary Table 1)** using LTR_STRUC (McCarthy et McDonald 2003). We preferred LTR_STRUC over other software since the latter still gives a few false detections of elements in plant genomes (R. Guyot, personal communication). Each set of predicted LTR-RTs for a given species was processed with Inpactor (Orozco-Arias et al. 2018) to classify elements into superfamilies (*Gypsy* or *Copia*) and sub-classify them into lineages according to the similarities of five amino acid reference domains (GAG, AP, RT, RNAseH, and INT). Once classified into superfamilies and lineages, the LTR-RTs of a genome are aligned against the elements of other genomes by pairs using BLASTn. The BLASTn bit-scores representing the number of matching and mis-matching residues (according to a given substitution matrix) between species (so called ‘all against all’) were displayed by a heatmap. The heatmap is organized according to the general phylogeny of the species used in this study (displayed **Supplementary Figure 1**) to appreciate both similarity and distribution of the elements analyzed (**Figure 1**).

**Figure 1.**
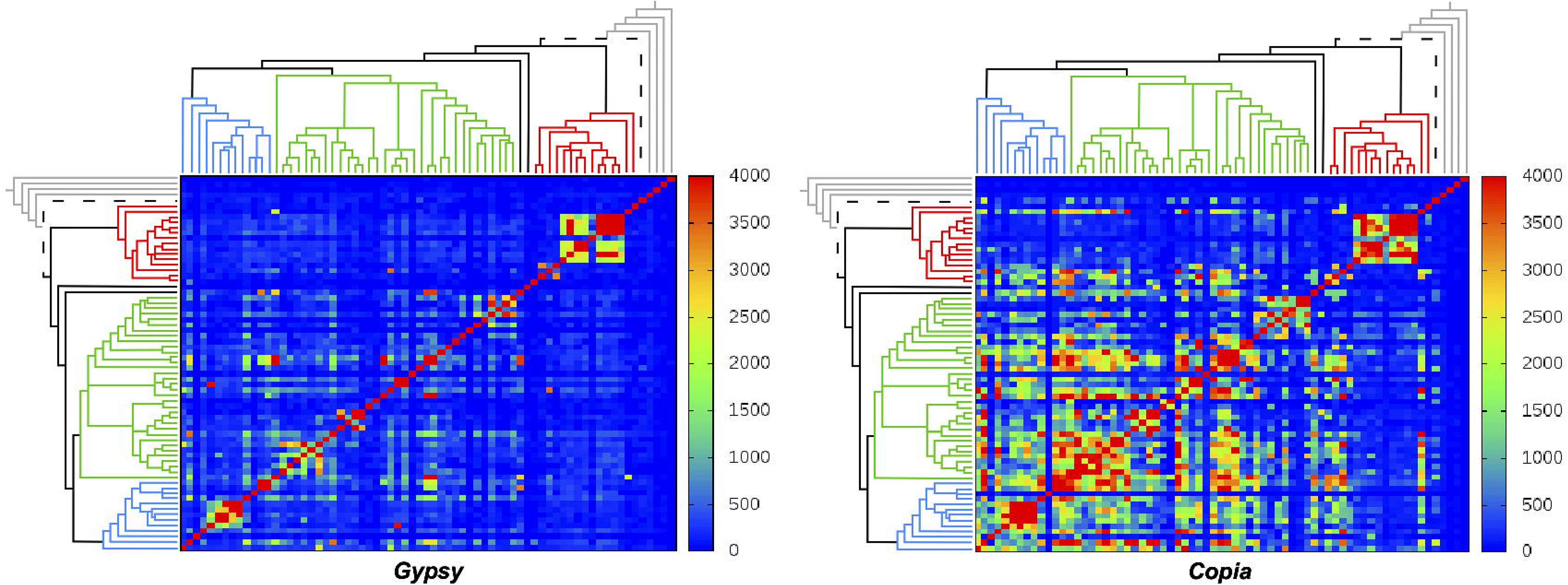
Representation of the similarity of *Gypsy* and *Copia* elements per genome across 69 plant species. Each *Gypsy* or *Copia* set of elements per genome is pairwise compared to the others using BLASTn. The level of similarity is represented by a heatmap of the BLAST scores (0 in blue: no similarity, to a maximum score of 4,000 in red). Species are organized in the matrix as shown in Sup. Fig. 1. A symbolic tree of the species was placed laterally and horizontally. Blue: Eudicots Asterids, Green: Eudicots Rosids, black: basal dicot species, red: Monocots, black dashed: Amborella and Grey: non-angiosperm. The diagonal represents the similarity of *Gypsy* and *Copia* elements of a species against itself.

For *Gypsy*, high pairwise BLASTn bit-scores were mainly restricted to monocots and to Brassicaceae, legumes and Solanaceae to a lesser extent. Some punctual high scores were also observed such as between cannabis and kiwi, castor bean and eggplant, or sacred lotus and clementine. For *Copia*, high scores were observed among all groups of species, except for *Amborella trichopoda, Utricularia gibba* (Asterids) and the non-angiosperm species (*Chlamydomonas reinhardtii, Physcomitrella patens, Selaginella moellendorffii, Picea abies*) (**Figure 1**). Interestingly, the distribution of the observed scores can be divided into three groups: dicotyledonous, sharing most of the high similarity across the group, monocotyledonous, with a high similarity also across them (mainly cereals), and orchids, Amborella and the non-angiosperm species, for which no significant pairwise scores were observed. Similar analyses were performed and confirmed with different upper BLASTn bit-score limits, ranging from 3,500 to 6,000 (**Supplementary Figure 2**).

In addition to superfamilies, LTR-RTs were sub-classified into lineages or groups of closely related lineages. Briefly, *Copia* lineages (i.e. Retrofit/Ale, Angela, Bianca, Oryco/Ivana, Tork/Tar/Ikeros, and SIRE) and *Gypsy* lineages (i.e. Athila, CRM, Del/Tekay, Galadriel, Reina, TAT) were identified according to the similarities in their internal amino acid domains (GAG, AP, INT, RT, RNAseH), as found in GyDB ((C. Llorens et al. 2011), http://gydb.org) and REXDB (Neumann et al. 2019) (**Supplementary Table 3**). The results (BLASTn score with an upper limit of 4,000) were plotted as previously, using a heatmap representation organized according to the phylogenetic order of the species.

For *Gypsy,* a clear patchy distribution of similarity and difference in the similarity pattern across the six lineages was observed (**Figure 2**). CRM appears as the *Gypsy* lineage with the highest scores. Breaks in the red diagonal (BLASTn analysis of the LTR-RTs set against itself) suggested that the lineage was not detected by LTR_STRUC and Inpactor. It is interesting to note that the cereal group (corresponding to the red lines in the phylogenetic tree representation) consistently showed a significant level of high scores across species and LTR-RT lineages. The Chlamyvir, Phygy and Selgy LTR-RT lineages were not found similar between any species (data not shown). For *Copia*, the similarity pattern differs considerably from that of *Gypsy* (**Figure 3**), although a patchy distribution of similarity is also evident. Similar to the superfamily analyses, high scores are observed for the lineages. The highest scores are unambiguously for the Tork/Tar/Ikeros, Retrofit/Ale, and Oryco/Ivana lineages. As with *Gypsy* lines, *Copia* lines of cereals have always shown strong pairwise similarities among cereal species. This indicates that the highest level of similarity is found among cereals species rather than with other angiosperms and non-angiosperm plant species. Finally, the Bryco, Lyco, Gymco and Osser lineages were not found similar between species (data not shown).

**Figure 2.**
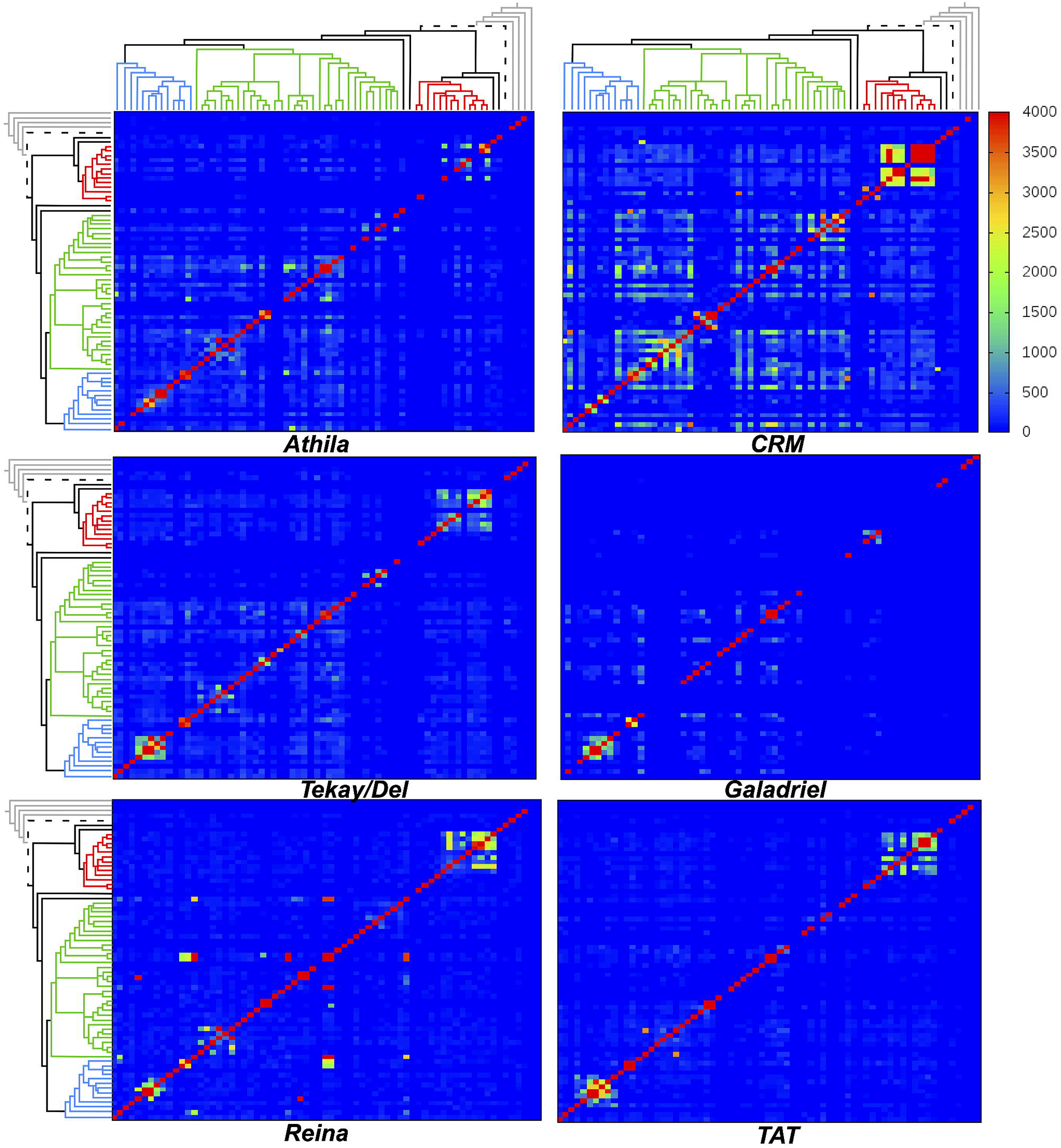
Representation of the levels of similarity of the *Gypsy* lineages elements per genome across 69 plant species. Each lineage (Athila, CRM, Tekay/Del, Galadriel, Reina and TAT) set of elements per genome is compared to the others using BLASTn. The level of similarity is represented by a heatmap of the BLASTn score (0 in blue: no similarity to a max BLAST score of 4,000 in red). Species are organized in the matrix as shown in Sup. Figure 1. The Chlamyvir, Phygy and Selgy lineages are not displayed.

**Figure 3.**
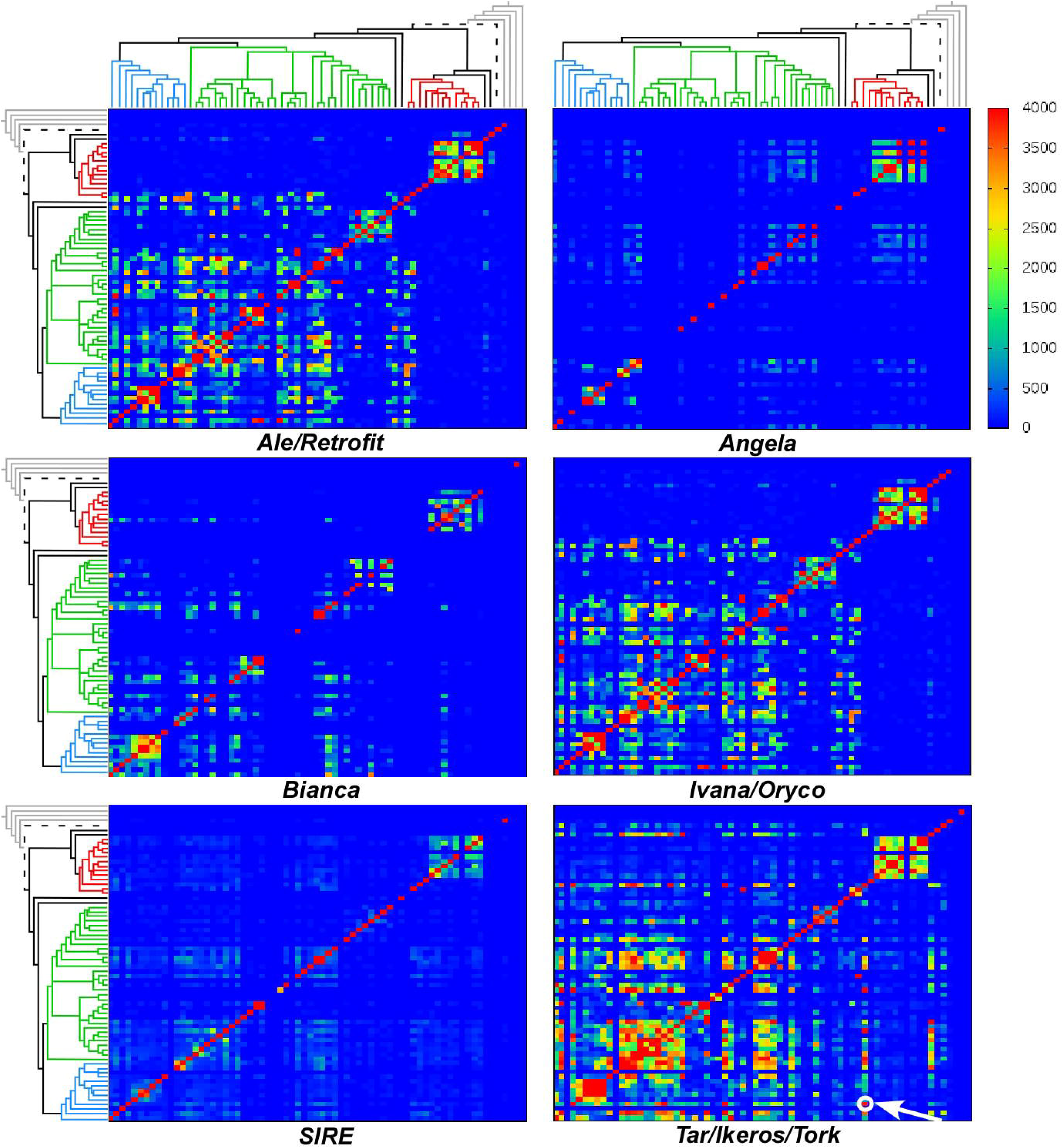
Representation of the similarity levels of Copia lineages per genome across 69 plant species. Each lineage (Bianca, Ivana/Oryco, Ale/Retrofit, SIRE and Tork/Tar/Ikeros) set of elements per genome is compared to the others using BLASTn. The level of similarity is represented by a heatmap of the BLASTn score (0 in blue: no similarity, to a max blast score of 4,000). Species are organized in the matrix as shown in Sup. Fig. 1. The Bryco, Lyco, Gymco and Osser lineages were not displayed. The white circle and arrow on the Tork/Tar/Ikeros panel indicates similarities between coffee and banana (Dias et al., 2015).

Among the 69 plant species analyzed here, orange (*C.sinensis*), eucalyptus (*E.grandis*) and prunus (*P. avium*) showed the highest number of highly similar elements with other species (**Figure 4A**). At the family level, most of the similarity was observed within the dicotyledonous family and little dicot/monocot similarity was noted at this stage of the analysis (BLASTn, score cutoff 4,000). Most of these similarities involved *Copia* lineages (Oryco/Ivana, Retrofit/Ale, Tork/Tar/Ikeros). This finding supports our previous observations using ‘heatmap’ representations (**Figure 4B and Supplementary Table 3**). The Oryco/Ivana, Retrofit/Ale, and Tork/Tar/Ikeros lineages are also the most similar elements between the different plant families (**Figure 4C**). In total, we noted 143 cases of high similarity, among them, 98 belong to three *Copia* lineages (Oryco/Ivana, Retrofit/Ale, Tork/Tar/Ikeros), representing 69% of the highly similar elements. Interestingly, we found that a previously reported detailed case of HTT between coffee and banana (Dias et al. 2015) was clearly displayed on the heatmap of similarity (**Figure 3**, white circle). Overall, these results indicate that there is a strong similarity between LTR-RTs involving mainly three lineages of *Copia*, detected in distant angiosperm species. Such similarity of LTR-RTs between distant species raises the question of whether some of them could be the result of horizontal transfers.

**Figure 4.**
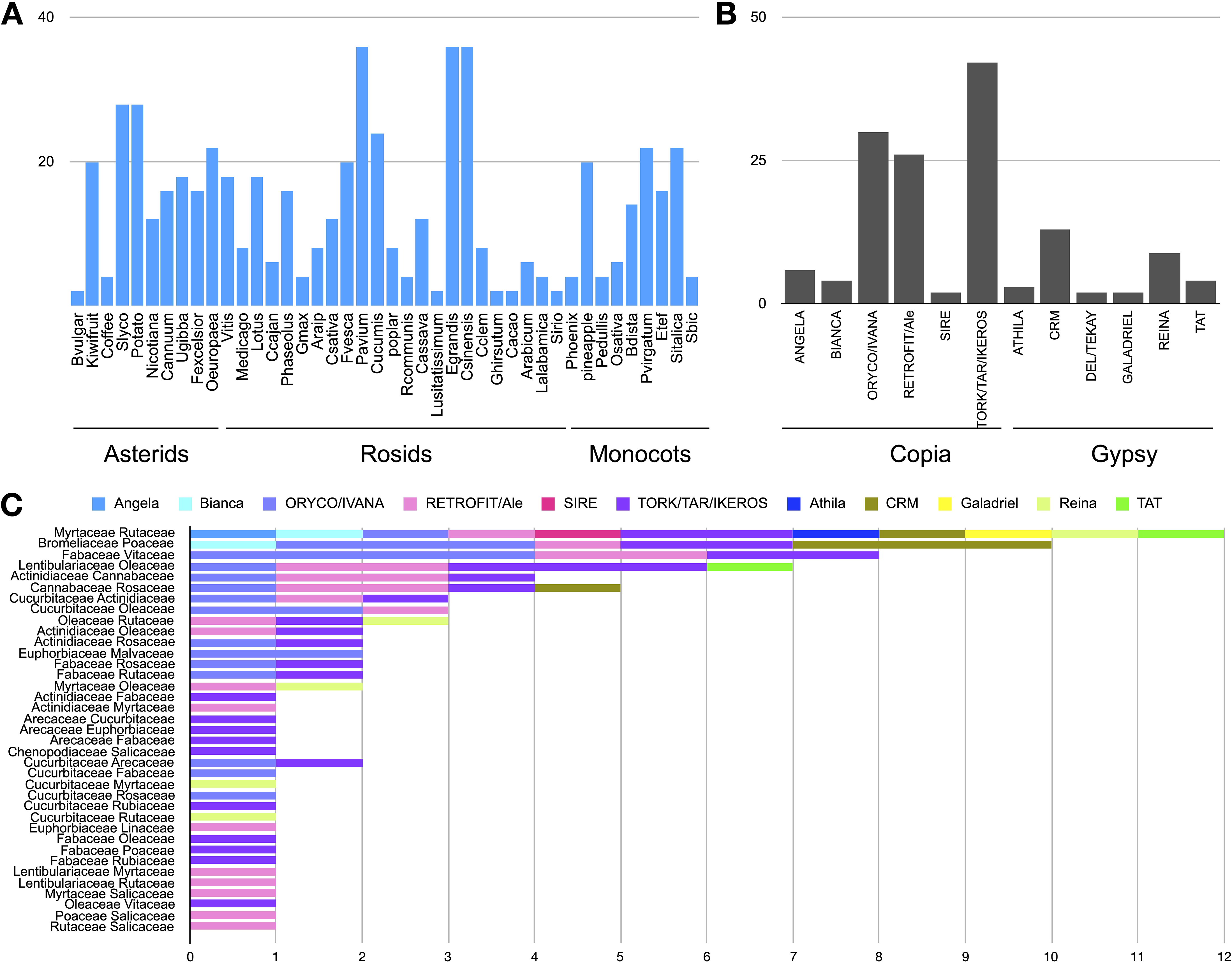
High similarity of LTR retrotransposons among 69 plant species. **A.** Number of LTR retrotransposons identified in one species and shared with another species. Species with no shared elements are not displayed. **B.** Number of shared LTR retrotransposon lineages found. *Gypsy* (Athila, CRM, Galadriel, Reina, TAT) and *Copia* (Angela, Bianca, Oryco/Ivana, Retrofit/Ale, SIRE, Tork/Tar/Ikeros) **C.** LTR retrotransposon lineages found highly similar only between different plant families.

### Detailed analyses of three randomly selected cases of LTR-RT high similarity

The high sequence similarities identified and the patchy distribution above are two of the criteria for identifying HT between species. To confirm the presence of potential HTs of LTR-RTs, other methods were applied on three different cases of high similarity identified different plant families. The three cases were randomly sampled. This small number if cases allow us to performed a rapid detailed analysis. We first analyzed the percentage of nucleotide identity of paired elements, since *K*s values may be inappropriate for distant transfers (El Baidouri et al. 2014). We analyzed the high similarity found between orchid (*Phalaenopsis equestris*, monocots) and cannabis (*Cannabis sativa*, Rosids; BLASTn score 4,276), between banana (*Musa acuminata*, monocots) and soybean (*Glycine max*, Rosids; BLASTn score 4,414), and between poplar (*Populus poplar*, Rosids) and coffee (*Coffea canephora*, Asteris; BLASTn score 4,597). The elements were first graphically aligned to estimate the degree of similarity across the full-length elements (coding and non-coding regions) and, then, they were pairwise aligned using nucmer (Kurtz et al. 2004) (**Figure 5**). All comparisons exhibited a high level of nucleotide identity between LTR-RTs, even in the non-coding regions (i.e. Long Terminal Repeats). This suggests that the high scores observed in the previous analysis indeed correspond to a very high nucleotide sequence similarity between the LTR-RTs. At the nucleotide level, orchid and cannabis elements show 75.2% of identity, while banana and soybean, and poplar and coffee exhibited 73.8% and 72.3% of shared residues, respectively. The similarity of the polyprotein genes (percentage of nucleotide identity) was also compared to the genome-wide sequence identity across all annotated genes for each genome pair analyzed (**Figure 5**). The peak values of the identity distribution between pairs of coding genes were lower than that of the conserved LTR retrotransposons. The phylogenetic distribution of these three elements was also studied. We extracted the RT domains of all elements recovered from the LTR_STRUC data set. RT classified as Tork/Tar/Ikeros (1,235 from 58 species) were aligned and used for phylogenetic analysis (**Figure 6**). Maximum Likelihood tree showed phylogenetic incongruences for the three elements studied. Tork/Tar/Ikeros elements from Orchid and Cannabis (**Figure 6, A**), Banana and Soybean (**B**) and Poplar and Coffee (**C**) clustered together showing a patchy distribution of these elements. Together with the phylogenetic incongruity, we concluded that HT might be one of the most probable mechanisms explaining the high BLASTn scores and high nucleotide similarity of the three Tork/Tar/Ikeros elements studied here.

**Figure 5.**
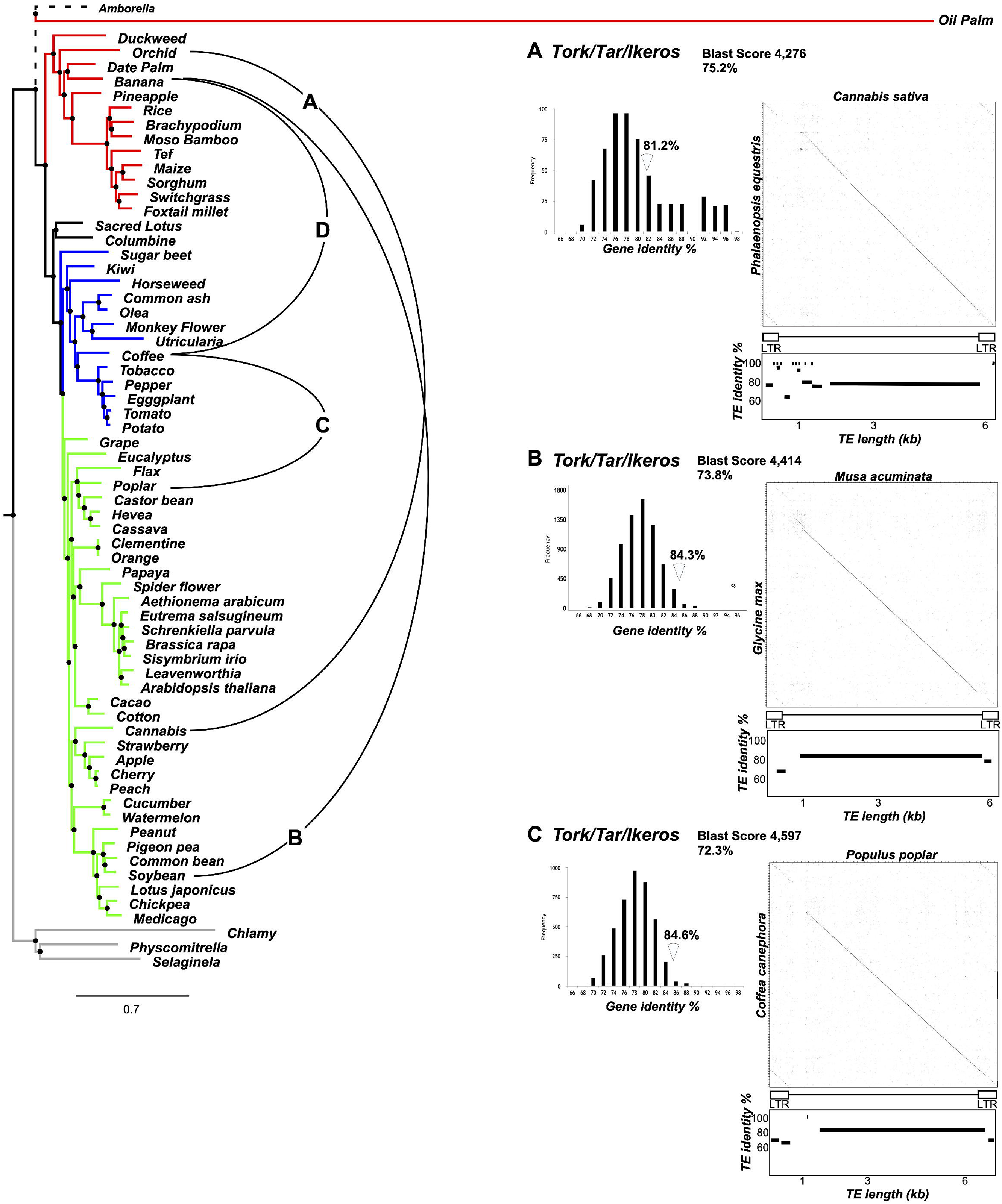
Detailed analysis of three potential cases of horizontal transfer identified in 69 sequenced plant genomes. The nuclear phylogenetic tree of the 69 plant genomes was computed using shared BUSCO genes. The cases of horizontal transfer analyzed are represented by colored lines connecting species and include the analysis of the nucleotide identity (%) of plant genes and elements, a dot-plot of conserved elements, and a graphical representation of the percentage of the identity (%) between shared elements. **A.** Analysis of HT between cannabis and orchid. **B**. Analysis of HT between banana and soybean. **C.** Analysis of HT between poplar and coffee. **D.** Relationships established in Dias et al., 2015.

**Figure 6.**
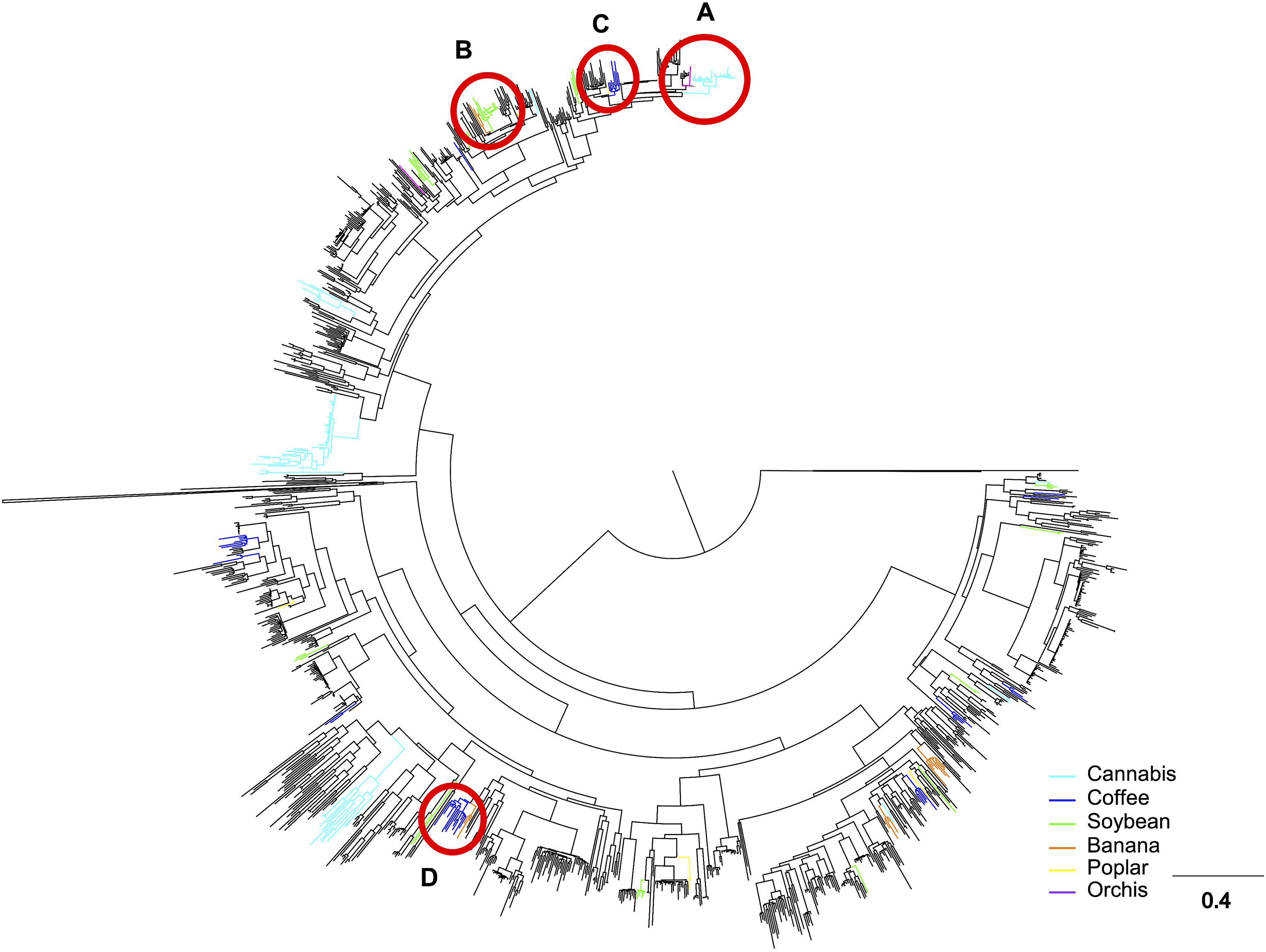
Phylogenetic tree of Tork RT domain extracted from 69 species LTR-RT prediction. Branches corresponding to Cannabis, Coffee, Soybean, Banana, poplar and Orchid are colored. Red circles indicate studied cases in Figure 5 (i.e., **A.** Cannabis and Orchid, **B.** Soybean and Banana, **C.** Coffee and Poplar) and **D.** illustrate a potential case of HT between Banana and Coffee (Dias et al., 2015).

### Reclassification of LTR-RT horizontally transferred

In 2014, using 40 different plant species, El Baidouri and coworkers (El Baidouri et al. 2014) found 32 new HTTs of LTR retrotransposons, yet the horizontally transferred HT elements were not classified at the lineage level. Thus, a classification was carried out using the same methodology as implemented in this study. It was found that only three HTT events included *Gypsy* elements (CRM: 2, Reina: 1) and the remaining 29 included *Copia* elements (Ale/Retrofit: 14, Bianca: 1, Ivana/Oryco: 7, Sire: 1, Tork/Tar/Ikeros: 6). Furthermore, the three most common lineages found in the previously proposed HTT events correspond to those found by this study (Ale/retrofit, Ivana/Oryco, and Tork/Tar/Ikeros) with the 84.3% of cases (i.e. 27 out of 32).

## DISCUSSION

The objective of the present study was to report and understand why some superfamilies and lineages of LTR retrotransposons displayed high similarities, even between distantly related plant species. Of course, such exceptional similarity strongly suggests potential cases of horizontal transfers since it is known that such transfers can happen between plants (El Baidouri et al. 2014). The strong similarity of two copies of an element between distant species is one of the required criteria to consider cases of horizontal transfers, together with the patchy distribution of the element across the phylogenetic tree of the species, and the phylogenetic incongruence between species and elements phylogenetic trees (Aubin, El Baidouri, et Panaud 2021).

Here, using 69 species, we show that there are many instances of very high similarity of *Copia* and *Gypsy* LTR-RTs between distantly related species, with a clear and patchy phylogenetic distribution, raising the question of whether these similarities could be considered as cases of horizontal transfers. Among the 143 high similarities observed, we analyzed in detail three of them, showing at the same time a high similarity, a patchy distribution and a phylogenetic incongruence, which corresponds to the three required criteria to consider horizontal transfer events. Alternative mechanisms to horizontal transfers must also be considered, like unequal substitution rates in TE sequences in different species (Silva et Kidwell 2000). For example, if paralogous copies of a same TE family in different lineages were vertically transmitted through speciation events, these copies could then be sampled in different species, leading to a TE phylogeny not matching the species tree (Loreto, Carareto, et Capy 2008). This scenario being possible mostly for related species, it can be excluded for some of the potential HTs detected in this study, involving very divergent plant species.

Although, we have to be careful with the hypothesis of HTs events for closely related plant species, the three potential HT cases studied in more details involve highly divergent species (monocot versus dicots, and Asterids versus Rosids), and the high similarity between the copies of each *Copia* element, including in their non-coding parts, is better explained by HT events than the latter hypotheses. Although we cannot rule out scenarios other than HT events for the set of elements found with high similarity in this study, these results provide basic evidence that may suggest that LTR-RT HTs may well be more frequent than expected in plants.

One of the key questions about HTs in plants is the identification of mechanisms or vectors able to carry TEs between different organisms. In insects, mite parasites, endosymbiotic bacteria, and viruses have been found to participate in the transfer of TEs (reviewed in (Gilbert et al. 2014; Wallau, Vieira, et Loreto 2018; Aubin, El Baidouri, et Panaud 2021)). In plants, strong suspicions are placed on viruses since they are able to infect a large range of different species, thus providing a strong argument in favor of their involvement in the transfer of host genetic material. Moreover, the ability of viruses to co-encapsulate host RNA, including LTR retrotransposons, has been recently demonstrated (Shrestha et al. 2018), suggesting the formation of active infectious viral particles.

Based on our analysis, SIRE (*Copia*) and Athila (*Gypsy*) show a very low level of interspecies similarity, although these lineages carry an envelope-like gene (*env,* allowing a retrovirus particle to leave the host cell) (Havecker, Gao, et Voytas 2004) and are thus considered potential endogenous retroviruses. This finding strongly suggests that the presence of an *env*-like gene is not the crucial factor involved in HTs of LTR-RT across plant species. This is clearly different from drosophila, where *Gypsy* elements are frequently found involved in HTs (Bartolomé, Bello, et Maside 2009), higher than *Copia* elements (Schaack, Gilbert, et Feschotte 2010). In particular, *Gypsy* in drosophila possess an *env*-like open reading frame and have been proposed as retrovirus-like particles able to infect other cells and organisms. In plants, different mechanisms allowing HTs might operate, and the presence of *env*-like genes in some LTR-RT lineages could represent an obstacle to successful HT events, contrary to what is observed in animal genomes.

Our analysis indicates that among *Copia* and *Gypsy* lineages classified so far in plants, only three related *Copia* lineages or groups of lineages, Ivana/Oryco, Ale/Retrofit and Tork/Tar/Ikeros are frequently found with high similarities and patchy distribution, and so potentially involved in HTs. This observation is supported by the reclassification of the 32 LTR-RT horizontally transferred in plants (El Baidouri et al. 2014) for which most of them (27) are *Copia* from Ivana/Oryco, Ale/Retrofit and Tork/Tar/Ikeros. Moreover, a re-analysis of the recent bibliography on plant HTs, indicates that *Copia* elements are clearly more frequently transferred than *Gypsy* (Dias et al. 2015; Roulin et al. 2007; 2009; Cheng et al. 2009; Huang et al. 2017; Hou et al. 2018; Park et al. 2021; Park, Christin, et Bennetzen 2021). Although classification data are not homogeneous, among the 175 cases of HT found in the literature, 65% of the LTR-RTs involved in HT are classified as *Copia*. When classification at the lineage level is available, the most transferred lineages are Tork and Ale/Retrofit (**Supplemental Table 4**). This observation is quite unexpected and raises important questions: what are the mechanisms allowing a high similarity between elements belonging to these 3 lineages of *Copia*? What would be the mechanisms of HT that would specifically select/favor these lineages?

Different tracks/leads should be pursued in the future to understand why some lineages are more conserved than others or more subject to horizontal transfers, such as their transcriptional and insertional activities and their copy number in the genomes.

## Acknowledgments

The authors acknowledge the IRD itrop HPC (South Green Platform) at IRD Montpellier for providing HPC resources that contributed to the research results reported in this paper. The authors thank the Universidad Autónoma de Manizales, Manizales, Colombia, for support under the project 752-115 and the LMI BIO-INCA for supporting Romain Guyot. We thank Moaine El Baidouri for his valuable comments on the Manuscript.

