## Supplementary material for "High nucleotide similarity of three *Copia* lineage LTR retrotransposons among plant genomes": Sup figure 1

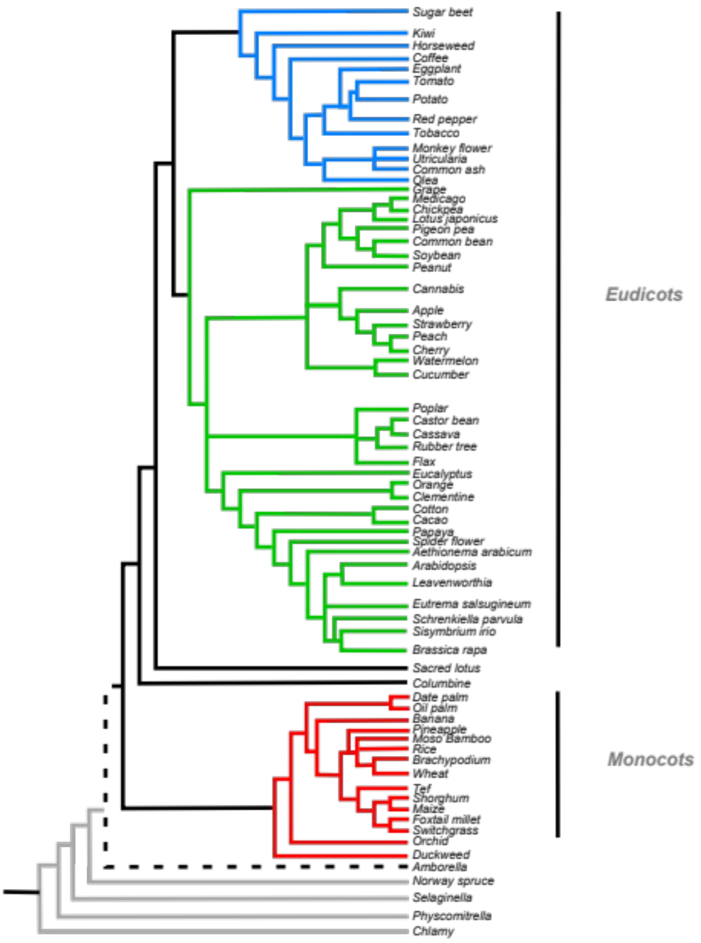


Sup. Figure 1. *Phylogenetic representation of the species used in the analysis. The scientific names are given in Supplementary Material 1. In grey: non-angiosperm species, in dashed black: Amborella, in red: monocots species, in green: Eudicots Rosids, and in blue: Eudicots Asterids.*
