## Supplementary material for "High nucleotide similarity of three *Copia* lineage LTR retrotransposons among plant genomes": Sup Figure 2

Supplementary Material 3

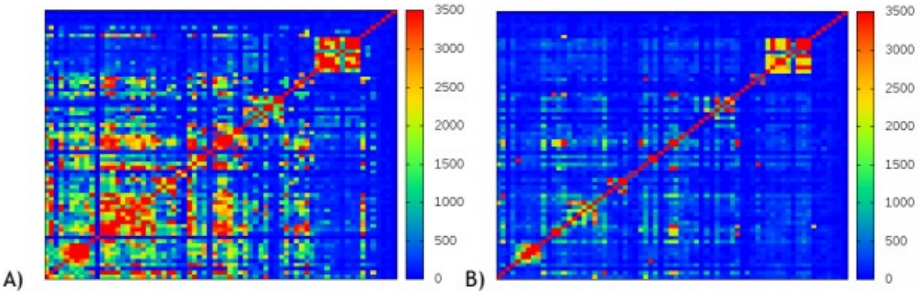


Heat maps of A) Copia and B) Gypsy using as max value 3,500

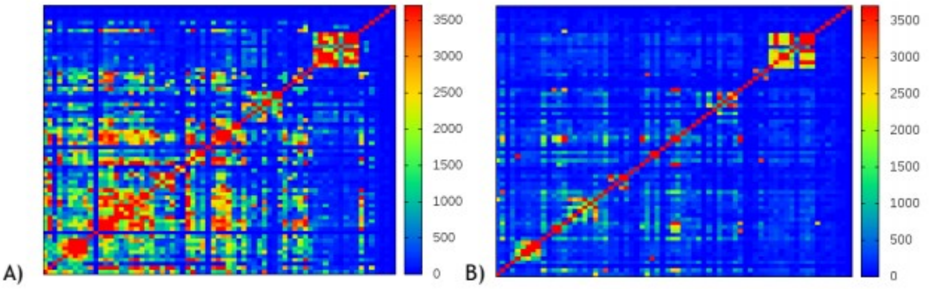


Heat maps of A) Copia and B) Gypsy using as max value 3,700

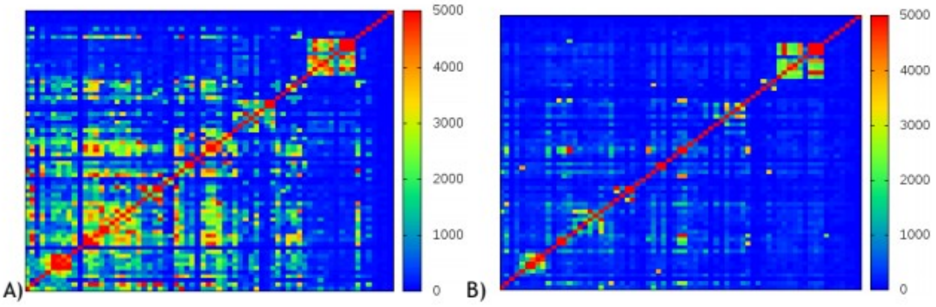


Heat maps of A) Copia and B) Gypsy using as max value 5,000

Sup. Figure 2.
