## Supplementary material for "High nucleotide similarity of three *Copia* lineage LTR retrotransposons among plant genomes": Sup Table 3

| Scientific Name | Common Name | Family | Genome size (Mb) | Source | URL |
|---|---|---|---|---|---|
| ***Amborella trichopoda*** | - | Amborellaceae | 685 | JGI | http://genome.jgi.doe.gov/pages/dynamicOrganismDownload.jsf?organism=Phytozome# (V12/ASSEMBLY/Atrichopoda_291_v1.0.fa.gz) |
| ***Elaeis oleifera*** | American oil Palm | Arecaceae | 1352 | NCBI | ftp://ftp.ncbi.nlm.nih.gov/genomes/all/GCA/000/441/515/GCA_000441515.1_EO8/GCA_000441515.1_EO8_genomic.fna.gz |
| ***Phoenix dactylifera*** | Date palm | Arecaceae | 547 | NCBI | ftp://ftp.ncbi.nlm.nih.gov/genomes/all/GCF/000/413/155/GCF_000413155.1_DPV01/GCF_000413155.1_DPV01_genomic.fna.gz |
| ***Arabidopsis thaliana*** | - | Brassicaceae | 116 | JGI | ftp://ftp.jgi-psf.org/pub/compgen/phytozome/v9.0/Athaliana/assembly/ |
| ***Brassica rapa*** | Chinese cabbage | Brassicaceae | 305 | JGI | ftp://ftp.jgi-psf.org/pub/compgen/phytozome/v9.0/Brapa/assembly/ |
| ***Ananas comosus*** | Pineapple | Bromeliaceae | 370 | JGI | http://genome.jgi.doe.gov/pages/dynamicOrganismDownload.jsf?organism=Acomosus# (V3.0/assembly/Acomosus_321_v3.fa.gz) |
| ***Carica papaya*** | papaya | Caricaceae | 332 | JGI | http://genome.jgi.doe.gov/pages/dynamicOrganismDownload.jsf?organism=Cpapaya# (ASGPBv0.4/assembly/Cpapaya_113_r.Dec2008.fa.gz) |
| ***Cucumis sativus*** | Cucumber | Cucurbitaceae | 365 | JGI | ftp://ftp.jgi-psf.org/pub/compgen/phytozome/v9.0/Csativus/assembly/ |
| ***Citrullus lanatus*** | watermelon | Cucurbitaceae | 313 | NCBI | ftp://ftp.ncbi.nlm.nih.gov/genomes/all/GCA/000/238/415/GCA_000238415.1_CiLa_1.0/GCA_000238415.1_CiLa_1.0_genomic.fna.gz |
| ***Manihot esculenta*** | Cassava | Euphorbiaceae | 564 | JGI | ftp://ftp.jgi-psf.org/pub/compgen/phytozome/v9.0/Mesculenta/assembly/ |
| ***Ricinus communis*** | Castor bean | Euphorbiaceae | 340 | JGI | http://genome.jgi.doe.gov/pages/dynamicOrganismDownload.jsf?organism=Rcommunis# (V0.1/assembly/Rcommunis_119_TIGR.0.1.fa.gz) |
| ***Cicer arietinum*** | chickpea | Fabaceae | 513 | NCBI | ftp://ftp.ncbi.nlm.nih.gov/genomes/all/GCF/000/331/145/GCF_000331145.1_ASM33114v1/GCF_000331145.1_ASM33114v1_genomic.fna.gz |
| ***Glycine max*** | soybean | Fabaceae | 939 | JGI | http://genome.jgi.doe.gov/pages/dynamicOrganismDownload.jsf?organism=Gmax# (V1.1/assembly/Gmax_189.fa.gz) |
| ***Lotus japonicus*** | Bird's-foot Trefoil | Fabaceae | 433 | Codon Usage Database | ftp://ftp.kazusa.or.jp/pub/lotus/lotus_r3.0/Lj3.0_pseudomol.fna.gz |
| ***Medicago truncatula*** | Barrel Medic | Fabaceae | 399 | JGI | ftp://ftp.jgi-psf.org/pub/compgen/phytozome/v9.0/Mtruncatula/assembly/ |
| ***Phaseolus vulgaris*** | common bean | Fabaceae | 518 | JGI | ftp://ftp.jgi-psf.org/pub/compgen/phytozome/v9.0/Pvulgaris/assembly/ |
| ***Utricularia gibba*** | humped bladderwort | Lentibulariaceae | 97 | NCBI | ftp://ftp.ncbi.nlm.nih.gov/genomes/all/GCA/002/189/035/GCA_002189035.1_U_gibba_v2/GCA_002189035.1_U_gibba_v2_genomic.fna.gz |
| ***Selaginella moellendorffii*** | - | Lycopodiophyta | 205 | JGI | http://genome.jgi.doe.gov/pages/dynamicOrganismDownload.jsf?organism=Smoellendorffii# (V1.0/assembly/Smoellendorffii_91_v1.fa.gz) |
| ***Theobroma cacao*** | Cacao tree | Malvaceae | 335 | JGI | ftp://ftp.jgi-psf.org/pub/compgen/phytozome/v9.0/Tcacao/assembly/ |
| ***Musa acuminata*** | Banana | Musaceae | 458 | JGI | http://genome.jgi.doe.gov/pages/dynamicOrganismDownload.jsf?organism=Phytozome# (v12/assembly/Macuminata_304_v1.fa.gz) |
| ***Mimulus guttatus*** | - | Phrymaceae | 302 | JGI | ftp://ftp.jgi-psf.org/pub/compgen/phytozome/v9.0/Mguttatus_v1.1/assembly/ |
| ***Brachypodium distachyon*** | purple false brome | Poaceae | 262 | JGI | ftp://ftp.jgi-psf.org/pub/compgen/phytozome/v9.0/Bdistachyon/assembly/ |
| ***Oryza sativa*** | rice | Poaceae | 364 | JGI | ftp://ftp.jgi-psf.org/pub/compgen/phytozome/v9.0/Osativa/assembly/ |
| ***Sorghum bicolor*** | - | Poaceae | 687 | JGI | ftp://ftp.jgi-psf.org/pub/compgen/phytozome/v9.0/Sbicolor_v1.4/assembly/ |
| ***Zea mays*** | Maize | Poaceae | 2068 | JGI | ftp://ftp.jgi-psf.org/pub/compgen/phytozome/v9.0/Zmays/assembly/ |
| ***Fragaria vesca*** | wild strawberry | Rosaceae | 201 | JGI | ftp://ftp.jgi-psf.org/pub/compgen/phytozome/v9.0/Fvesca/assembly/ |
| ***Malus domestica*** | apple | Rosaceae | 859 | JGI | ftp://ftp.jgi-psf.org/pub/compgen/phytozome/v9.0/Mdomestica/assembly/ |
| ***Prunus persica*** | peach | Rosaceae | 219 | JGI | ftp://ftp.jgi-psf.org/pub/compgen/phytozome/v9.0/Ppersica/assembly/ |
| ***Coffea canephora*** | robusta coffee | Rubiaceae | 552 | Coffee Genome Hub | http://coffee-genome.org/coffeacanephora |
| ***Citrus clementina*** | Clementine | Rutaceae | 291 | JGI | ftp://ftp.jgi-psf.org/pub/compgen/phytozome/v9.0/Cclementina/assembly/ |
| ***Populus trychocarpa*** | poplar | Salicaceae | 419 | JGI | ftp://ftp.jgi-psf.org/pub/compgen/phytozome/v9.0/Ptrichocarpa/assembly/ |
| ***Nicotiana sylvestris*** | Tobacco plant | Solanaceae | 2171 | NCBI | ftp://ftp.ncbi.nlm.nih.gov/genomes/all/GCF/000/393/655/GCF_000393655.1_Nsyl/GCF_000393655.1_Nsyl_genomic.fna.gz |
| ***Solanum tuberosum*** | potato | Solanaceae | 747 | JGI | http://genome.jgi.doe.gov/pages/dynamicOrganismDownload.jsf?organism=Stuberosum# (V4.03/assembly/Stuberosum_448_v4.03.fa.gz) |
| ***Solanum lycopersicum*** | tomato | Solanaceae | 796 | JGI | ftp://ftp.jgi-psf.org/pub/compgen/phytozome/v9.0/Slycopersicum/assembly/ |
| ***Vitis vinifera*** | grape | Vitaceae | 471 | JGI | ftp://ftp.jgi-psf.org/pub/compgen/phytozome/v9.0/Vvinifera/assembly/ |
| ***Beta vulgaris*** | sugar beet | Chenopodiaceae | 541 | The Beta Bulgaris Resources | http://bvseq.molgen.mpg.de/Genome/Download/RefBeet-1.2/RefBeet-1.2.fna.gz |
| ***Eucalyptus grandis*** | Rose gum | Myrtaceae | 667 | JGI | http://genome.jgi.doe.gov/pages/dynamicOrganismDownload.jsf?organism=Phytozome# (Egrandis_v297_v2.0.fa.gz) |
| ***Prunus avium*** | sweet cherry | Rosaceae | 362 | MainLab Bioinformatics | ftp://ftp.bioinfo.wsu.edu/species/Prunus_avium/Prunus_avium-genome.v1.0.a1/assembly/Prunus_avium_v1.0.a1_pseudomolecule.fasta.gz |
| ***Capsicum annuum*** | Pepper | Solanaceae | 2826 | NCBI | ftp://ftp.ncbi.nlm.nih.gov/genomes/all/GCF/000/710/875/GCF_000710875.1_Pepper_Zunla_1_Ref_v1.0/GCF_000710875.1_Pepper_Zunla_1_Ref_v1.0_genomic.fna.gz |
| ***Panicum virgatum*** | switchgrass | Poaceae | 1638 | JGI | http://genome.jgi.doe.gov/pages/dynamicOrganismDownload.jsf?organism=Phytozome# (Pvirgatum_273_v1.0.fa.gz) |
| ***Triticum aestivum*** | bread wheat | Poaceae | 13005 | Ensambl Genomes | ftp://ftp.ensemblgenomes.org/pub/plants/release-36/fasta/triticum_aestivum/dna/Triticum_aestivum.TGACv1.dna.toplevel.fa.gz |
| ***Phalaenopsis equestris*** | Moth orchid | Orchidaceae | 1044 | CoGe | https://genomevolution.org/coge/GenomeInfo.pl?gid=25065 |
| ***Gossypium hirsutum*** | Cotton | Malvaceae | 2427 | MainLab Bioinformatics | ftp://ftp.bioinfo.wsu.edu/species/Gossypium_hirsutum/NAU-NBI_G.hirsutum_AD1genome/assembly/NBI_Gossypium_hirsutum_v1.1.fa.gz |
| ***Actinidia deliciosa*** | Kiwifruit | Actinidiaceae | 602 | Fei Lab BTI | ftp://bioinfo.bti.cornell.edu/pub/kiwifruit/Kiwifruit_pseudomolecule.fa.gz |
| ***Chlamydomonas reinhardtii*** | Green algae | Chlamydomonadaceae | 107 | JGI | http://genome.jgi.doe.gov/pages/dynamicOrganismDownload.jsf?organism=Creinhardtii# (V5.5/Assembly/Creinhardtii_281_v5.0.fa.gz) |

| | | | | | |
|---|---|---|---|---|---|
| ***Hevea brasiliensis*** | rubber tree | Euphorbiaceae | 1321 | NCBI | ftp://ftp.ncbi.nlm.nih.gov/genomes/all/GCA/001/654/055/GCA_001654055.1_ASM165405v1/GCA_001654055.1_ASM165405v1_genomic.fna.gz |
| ***Olea europaea*** | Olive tree | Oleaceae | 1270 | CNAG | http://denovo.cnag.cat/genomes/olive/download/Oe6/Oe6.scaffolds.fa.gz |
| ***Erigeron canadensis*** | Horseweed | Asteraceae | 316 | NCBI | ftp://ftp.ncbi.nlm.nih.gov/genomes/all/GCA/000/775/935/GCA_000775935.1_ASM77593v1/GCA_000775935.1_ASM77593v1_genomic.fna.gz |
| ***Solanum melongena*** | Eggplant | Solanaceae | 807 | NCBI | ftp://ftp.ncbi.nlm.nih.gov/genomes/all/GCA/000/787/875/GCA_000787875.1_SME_r2.5.1/GCA_000787875.1_SME_r2.5.1_genomic.fna.gz |
| ***Fraxinus excelsior*** | Common Ash | Oleaceae | 846 | NCBI | ftp://ftp.ncbi.nlm.nih.gov/genomes/all/GCA/900/149/125/GCA_900149125.1_BATG-0.5/GCA_900149125.1_BATG-0.5_genomic.fna.gz |
| ***Cajanus cajan*** | Pigeon Pea | Fabaceae | 576 | NCBI | ftp://ftp.ncbi.nlm.nih.gov/genomes/all/GCF/000/340/665/GCF_000340665.1_C.cajan_V1.0/GCF_000340665.1_C.cajan_V1.0_genomic.fna.gz |
| ***Arachis ipaensis*** | Peanut | Fabaceae | 1290 | PeanutBase | https://peanutbase.org/files/genomes/Arachis_ipaensis/assembly/Araip_v1.0.tar.gz |
| ***Cannabis Sativa*** | Cannabis | Cannabaceae | 566 | NCBI | ftp://ftp.ncbi.nlm.nih.gov/genomes/all/GCA/001/865/755/GCA_001865755.1_ASM186575v1/GCA_001865755.1_ASM186575v1_genomic.fna.gz |
| ***Linum usitatissimum*** | Flax | Linaceae | 277 | NCBI | ftp://ftp.ncbi.nlm.nih.gov/genomes/all/GCA/000/224/295/GCA_000224295.1_LinUsi_v1.1/GCA_000224295.1_LinUsi_v1.1_genomic.fna.gz |
| ***Citrus sinensis*** | Orange | Rutaceae | 317 | NCBI | ftp://ftp.ncbi.nlm.nih.gov/genomes/all/GCF/000/317/415/GCF_000317415.1_Csi_valencia_1.0/GCF_000317415.1_Csi_valencia_1.0_genomic.fna.gz |
| ***Tarenaya hassleriana*** | Spider Flower | Cleomaceae | 242 | NCBI | ftp://ftp.ncbi.nlm.nih.gov/genomes/all/GCF/000/463/585/GCF_000463585.1_ASM46358v1/GCF_000463585.1_ASM46358v1_genomic.fna.gz |
| ***Aethionema arabicum*** | - | Brassicaceae | 187 | NCBI | ftp://ftp.ncbi.nlm.nih.gov/genomes/all/GCA/000/411/095/GCA_000411095.1_VEGI_AA_v_1.0/GCA_000411095.1_VEGI_AA_v_1.0_genomic.fna.gz |
| ***Leavenworthia alabamica*** | - | Brassicaceae | 168 | NCBI | ftp://ftp.ncbi.nlm.nih.gov/genomes/all/GCA/000/411/055/GCA_000411055.1_VEGI_LA_v_1.0/GCA_000411055.1_VEGI_LA_v_1.0_genomic.fna.gz |
| ***Eutrema salsugineum*** | - | Brassicaceae | 234 | NCBI | ftp://ftp.ncbi.nlm.nih.gov/genomes/all/GCF/000/478/725/GCF_000478725.1_Eutsalg1_0/GCF_000478725.1_Eutsalg1_0_genomic.fna.gz |
| ***Schrenkiella parvula*** | - | Brassicaceae | 132 | NCBI | ftp://ftp.ncbi.nlm.nih.gov/genomes/all/GCA/000/218/505/GCA_000218505.1_Eutrema_parvulum_v01/GCA_000218505.1_Eutrema_parvulum_v01_genomic.fna.gz |
| ***Sisymbrium irio*** | - | Brassicaceae | 239 | NCBI | ftp://ftp.ncbi.nlm.nih.gov/genomes/all/GCA/000/411/075/GCA_000411075.1_VEGI_SI_v_1.0/GCA_000411075.1_VEGI_SI_v_1.0_genomic.fna.gz |
| ***Nelumbo nucifera*** | Sacred Lotus | Nelumbonaceae | 777 | NCBI | ftp://ftp.ncbi.nlm.nih.gov/genomes/all/GCF/000/365/185/GCF_000365185.1_Chinese_Lotus_1.1/GCF_000365185.1_Chinese_Lotus_1.1_genomic.fna.gz |
| ***Aquilegia coerulea*** | Columbine | Ranunculaceae | 295 | JGI | http://genome.jgi.doe.gov/pages/dynamicOrganismDownload.jsf?organism=Acoerulea# (V3.1/assembly/Acoerulea_322_v3.fa.gz) |
| ***Phyllostachys edulis*** | Moso bamboo | Poaceae | 1987 | ICBR | http://202.127.18.221/bamboo/files/P_heterocycla_v1.0.Scaffolds.gz |
| ***Eragrostis tef*** | tef | Poaceae | 587 | NCBI | ftp://ftp.ncbi.nlm.nih.gov/genomes/all/GCA/000/970/635/GCA_000970635.1_ASM97063v1/GCA_000970635.1_ASM97063v1_genomic.fna.gz |
| ***Setaria italica*** | Foxtail Millet | Poaceae | 391 | NCBI | ftp://ftp.ncbi.nlm.nih.gov/genomes/all/GCF/000/263/155/GCF_000263155.2_Setaria_italica_v2.0/GCF_000263155.2_Setaria_italica_v2.0_genomic.fna.gz |
| ***Spirodela polyrhiza*** | Duckweed | Araceae | 131 | NCBI | ftp://ftp.ncbi.nlm.nih.gov/genomes/all/GCA/001/981/405/GCA_001981405.1_ASM198140v1/GCA_001981405.1_ASM198140v1_genomic.fna.gz |
| ***Picea abies*** | norway Spruce | Pinaceae | 11981 | The Plant Genome Integrative Explorer | ftp://plantgenie.org/Data/ConGenIE/Picea_abies/v1.0/FASTA/GenomeAssemblies/Pabies1.0-genome.fa.gz |
| ***Physcomitrella patens*** | physcomitrella | Funariaceae | 461 | NCBI | ftp://ftp.ncbi.nlm.nih.gov/genomes/all/GCF/000/002/425/GCF_000002425.3_V1.1/GCF_000002425.3_V1.1_genomic.fna.gz |

| Family | Species used |
|---|---|
| Actinidiaceae | 1 |
| Amborellaceae | 1 |
| Araceae | 1 |
| Arecaceae | 2 |
| Asteraceae | 1 |
| Brassicaceae | 7 |
| Bromeliaceae | 1 |
| Cannabaceae | 1 |
| Caricaceae | 1 |
| Chenopodiaceae | 1 |
| Chlamydomonadaceae | 1 |
| Cleomaceae | 1 |
| Cucurbitaceae | 2 |
| Euphorbiaceae | 3 |
| Fabaceae | 7 |
| Funariaceae | 1 |
| Lentibulariaceae | 1 |
| Linaceae | 1 |
| Lycopodiophyta | 1 |
| Malvaceae | 2 |
| Musaceae | 1 |
| Myrtaceae | 1 |
| Nelumbonaceae | 1 |
| Oleaceae | 2 |
| Orchidaceae | 1 |
| Phrymaceae | 1 |
| Pinaceae | 1 |
| Poaceae | 9 |
| Ranunculaceae | 1 |
| Rosaceae | 4 |
| Rubiaceae | 1 |
| Rutaceae | 2 |
| Salicaceae | 1 |
| Solanaceae | 5 |
| Vitaceae | 1 |

Sup. Table 1

| Species | Number of predicted LTR-RTs |
|---|---|
| ***Bvulgar*** | 2070 |
| ***Kiwifruit*** | 1190 |
| ***Ecanadensis*** | 97 |
| ***Coffee*** | 1743 |
| ***Smelongena*** | 410 |
| ***Slyco*** | 1203 |
| ***Potato*** | 1217 |
| ***Nicotiana*** | 1434 |
| ***Cannuum*** | 1350 |
| ***Mgut*** | 2184 |
| ***Ugibba*** | 39 |
| ***Fexcelsior*** | 891 |
| ***Oeuropaea*** | 3787 |
| ***Vitis*** | 1372 |
| ***Medicago*** | 867 |
| ***Cicer*** | 726 |
| ***Lotus*** | 785 |
| ***Ccajan*** | 1390 |
| ***Phaseolus*** | 985 |
| ***Gmax*** | 3129 |
| ***Araip*** | 2606 |
| ***Csativa*** | 6733 |
| ***Malus*** | 1018 |
| ***Fvesca*** | 368 |
| ***Pavium*** | 480 |
| ***PrunusP*** | 1713 |
| ***watermelon*** | 118 |
| ***Cucumis*** | 52 |
| ***poplar*** | 368 |
| ***Rcommunis*** | 1066 |
| ***Cassava*** | 459 |
| ***Hbrasiliensis*** | 1729 |
| ***Lusitatissimum*** | 203 |
| ***Egrandis*** | 1549 |
| ***Csinensis*** | 651 |
| ***Cclem*** | 1838 |
| ***Ghirsutum*** | 1738 |
| ***Cacao*** | 823 |
| ***papa*** | 201 |
| ***Thassleriana*** | 461 |
| ***Arabicum*** | 90 |
| ***Ath*** | 195 |
| ***Lalabamica*** | 75 |
| ***Eusalsugineum*** | 1553 |
| ***Sparvula*** | 94 |
| ***Sirio*** | 148 |
| ***Brapa*** | 412 |
| ***Nnucifera*** | 1109 |
| ***Acoerulea*** | 84 |
| ***Phoenix*** | 72 |
| ***oilpalm*** | 374 |
| ***banana*** | 532 |
| ***pineapple*** | 187 |
| ***Pedullis*** | 1145 |
| ***Osativa*** | 2358 |
| ***Bdista*** | 759 |
| ***Taestivum*** | 9471 |
| ***Pvirgatum*** | 897 |
| ***Etef*** | 241 |
| ***Sitalica*** | 2257 |
| ***Sbic*** | 1754 |
| ***Zmays*** | 6546 |
| ***Pequestris*** | 331 |
| ***Spolyrhiza*** | 104 |
| ***AmTr*** | 364 |
| ***Pabies*** | 1348 |
| ***Selag*** | 1335 |
| ***Ppatens*** | 717 |
| ***Creinhardtii*** | 67 |

Sup Table 3 A

| Specie | Retrofit | Ale | Alesia | Angela | Bianca | Bryco | Lyco | Gymco | Ikeros | Ivana | Oryco | Osser | SIRE | TAR | Tork | No Lineage | Total Copia | CRM | DEL | Galadriel | Reina | Tekay | Athila | Tat | Phygy | Selgy | No Lineage | Total Gypsy |
|---|---|---|---|---|---|---|---|---|---|---|---|---|---|---|---|---|---|---|---|---|---|---|---|---|---|---|---|---|
| Bvulgar | 398 | 92 | 0 | 15 | 22 | 0 | 0 | 0 | 0 | 0 | 34 | 0 | 46 | 0 | 90 | 32 | 729 | 108 | 55 | 0 | 108 | 0 | 32 | 405 | 0 | 0 | 1 | 709 |
| Kiwifruit | 212 | 0 | 0 | 2 | 1 | 0 | 0 | 0 | 2 | 0 | 28 | 0 | 19 | 9 | 98 | 77 | 448 | 58 | 47 | 7 | 24 | 0 | 34 | 111 | 0 | 0 | 3 | 284 |
| Ecanadensis | 23 | 1 | 0 | 0 | 1 | 0 | 0 | 0 | 0 | 0 | 5 | 0 | 3 | 0 | 0 | 3 | 36 | 2 | 1 | 0 | 7 | 0 | 0 | 3 | 0 | 0 | 0 | 13 |
| Coffee | 113 | 0 | 0 | 3 | 66 | 0 | 0 | 0 | 0 | 0 | 31 | 0 | 106 | 6 | 128 | 31 | 484 | 76 | 101 | 7 | 115 | 0 | 93 | 178 | 0 | 0 | 3 | 573 |
| Smelongena | 54 | 1 | 1 | 0 | 4 | 0 | 1 | 0 | 1 | 0 | 4 | 0 | 17 | 5 | 15 | 12 | 115 | 10 | 74 | 10 | 4 | 1 | 8 | 3 | 0 | 0 | 3 | 113 |
| Slyco | 187 | 1 | 0 | 1 | 38 | 0 | 1 | 1 | 4 | 0 | 45 | 0 | 67 | 22 | 166 | 48 | 581 | 4 | 225 | 34 | 15 | 3 | 94 | 29 | 0 | 0 | 1 | 405 |
| Potato | 243 | 2 | 1 | 2 | 15 | 0 | 6 | 0 | 2 | 0 | 57 | 0 | 32 | 7 | 27 | 42 | 436 | 5 | 155 | 9 | 36 | 1 | 57 | 9 | 0 | 0 | 1 | 273 |
| Nicotiana | 97 | 0 | 0 | 1 | 18 | 0 | 0 | 0 | 1 | 0 | 5 | 0 | 89 | 2 | 64 | 17 | 294 | 2 | 221 | 18 | 21 | 1 | 107 | 239 | 0 | 0 | 9 | 618 |
| Cannuum | 103 | 0 | 0 | 1 | 6 | 0 | 0 | 0 | 2 | 2 | 27 | 0 | 17 | 15 | 43 | 40 | 256 | 24 | 252 | 4 | 33 | 1 | 122 | 23 | 0 | 0 | 1 | 460 |
| Mgut | 51 | 7 | 0 | 295 | 67 | 0 | 0 | 0 | 0 | 0 | 36 | 0 | 193 | 4 | 76 | 57 | 786 | 39 | 57 | 20 | 42 | 0 | 111 | 166 | 0 | 0 | 6 | 441 |
| Ugibba | 4 | 0 | 0 | 0 | 0 | 0 | 0 | 0 | 0 | 0 | 7 | 0 | 0 | 0 | 5 | 1 | 17 | 3 | 0 | 0 | 2 | 0 | 2 | 9 | 0 | 0 | 0 | 16 |
| Fexcelsior | 153 | 1 | 1 | 1 | 1 | 0 | 0 | 1 | 6 | 2 | 39 | 0 | 16 | 13 | 108 | 77 | 419 | 30 | 5 | 4 | 26 | 0 | 9 | 77 | 0 | 0 | 1 | 152 |
| Oeuropaea | 140 | 0 | 0 | 50 | 6 | 0 | 0 | 0 | 2 | 1 | 94 | 0 | 36 | 57 | 515 | 132 | 1033 | 301 | 96 | 8 | 73 | 0 | 83 | 537 | 0 | 0 | 9 | 1107 |
| Vitis | 401 | 0 | 0 | 12 | 39 | 0 | 0 | 0 | 0 | 1 | 60 | 0 | 19 | 6 | 169 | 111 | 818 | 24 | 8 | 25 | 27 | 0 | 132 | 109 | 0 | 0 | 0 | 325 |
| Medicago | 42 | 5 | 0 | 6 | 7 | 0 | 0 | 0 | 10 | 1 | 40 | 0 | 119 | 7 | 55 | 9 | 301 | 10 | 69 | 0 | 16 | 0 | 7 | 211 | 0 | 0 | 19 | 332 |
| Cicer | 142 | 14 | 13 | 0 | 0 | 0 | 0 | 0 | 0 | 3 | 43 | 0 | 42 | 1 | 86 | 34 | 378 | 3 | 6 | 0 | 33 | 0 | 7 | 1 | 0 | 0 | 0 | 50 |
| Lotus | 35 | 18 | 0 | 0 | 13 | 0 | 0 | 0 | 0 | 0 | 28 | 0 | 134 | 1 | 23 | 16 | 268 | 2 | 27 | 0 | 81 | 1 | 5 | 156 | 0 | 0 | 3 | 275 |
| Ccajan | 77 | 2 | 0 | 0 | 16 | 0 | 0 | 1 | 0 | 2 | 89 | 0 | 24 | 0 | 57 | 42 | 310 | 70 | 66 | 0 | 90 | 0 | 26 | 352 | 0 | 0 | 4 | 608 |
| Phaseolus | 29 | 0 | 0 | 0 | 0 | 0 | 0 | 0 | 1 | 1 | 55 | 0 | 69 | 1 | 35 | 18 | 209 | 77 | 6 | 0 | 33 | 0 | 2 | 274 | 0 | 0 | 5 | 397 |
| Gmax | 43 | 9 | 3 | 0 | 0 | 0 | 0 | 0 | 25 | 0 | 186 | 0 | 499 | 1 | 181 | 37 | 984 | 518 | 31 | 0 | 237 | 0 | 260 | 295 | 0 | 0 | 46 | 1387 |
| Araip | 108 | 0 | 0 | 0 | 40 | 0 | 0 | 0 | 1 | 0 | 14 | 0 | 64 | 0 | 51 | 16 | 294 | 5 | 23 | 7 | 23 | 0 | 369 | 582 | 0 | 0 | 16 | 1025 |
| Csativa | 274 | 3 | 1 | 443 | 9 | 0 | 0 | 0 | 19 | 1 | 10 | 0 | 315 | 1 | 439 | 91 | 1606 | 18 | 1414 | 14 | 23 | 0 | 138 | 453 | 0 | 0 | 8 | 2068 |
| Malus | 135 | 9 | 0 | 0 | 61 | 0 | 0 | 0 | 3 | 1 | 64 | 0 | 1 | 2 | 33 | 75 | 384 | 5 | 5 | 1 | 38 | 0 | 2 | 37 | 0 | 0 | 3 | 91 |
| Fvesca | 31 | 3 | 0 | 1 | 65 | 0 | 0 | 0 | 0 | 1 | 8 | 0 | 4 | 2 | 17 | 17 | 149 | 9 | 5 | 5 | 9 | 0 | 1 | 33 | 0 | 0 | 0 | 62 |
| Pavium | 58 | 7 | 0 | 0 | 11 | 0 | 0 | 0 | 0 | 0 | 41 | 0 | 6 | 0 | 23 | 55 | 201 | 9 | 1 | 0 | 5 | 0 | 1 | 16 | 0 | 0 | 0 | 32 |
| PrunusP | 149 | 9 | 0 | 0 | 70 | 0 | 0 | 0 | 0 | 0 | 176 | 0 | 18 | 7 | 62 | 155 | 646 | 34 | 15 | 2 | 8 | 0 | 38 | 219 | 0 | 0 | 0 | 316 |
| watermelon | 22 | 0 | 0 | 1 | 0 | 0 | 0 | 0 | 0 | 0 | 5 | 0 | 0 | 2 | 9 | 11 | 50 | 8 | 4 | 9 | 16 | 0 | 0 | 1 | 0 | 0 | 0 | 38 |
| Cucumis | 9 | 0 | 0 | 0 | 0 | 0 | 0 | 0 | 0 | 0 | 0 | 0 | 0 | 0 | 2 | 0 | 11 | 4 | 1 | 3 | 20 | 0 | 0 | 0 | 0 | 0 | 0 | 28 |
| poplar | 95 | 1 | 0 | 1 | 0 | 0 | 0 | 0 | 0 | 3 | 42 | 0 | 1 | 0 | 12 | 39 | 194 | 22 | 4 | 1 | 35 | 0 | 20 | 5 | 0 | 0 | 0 | 87 |
| Rcommunis | 206 | 1 | 0 | 0 | 0 | 0 | 0 | 0 | 0 | 3 | 62 | 0 | 4 | 0 | 60 | 84 | 420 | 110 | 42 | 45 | 12 | 0 | 63 | 4 | 0 | 0 | 1 | 277 |
| Cassava | 103 | 1 | 0 | 1 | 0 | 0 | 0 | 0 | 0 | 0 | 15 | 0 | 0 | 0 | 6 | 32 | 158 | 24 | 76 | 6 | 19 | 0 | 9 | 20 | 0 | 0 | 2 | 156 |
| Hbrasiliensis | 233 | 2 | 2 | 165 | 1 | 0 | 0 | 0 | 0 | 0 | 29 | 0 | 28 | 0 | 20 | 64 | 544 | 45 | 592 | 3 | 23 | 0 | 69 | 33 | 0 | 0 | 3 | 768 |
| Lusitatissimum | 68 | 0 | 0 | 0 | 0 | 0 | 0 | 0 | 0 | 2 | 37 | 0 | 9 | 0 | 18 | 21 | 155 | 1 | 3 | 0 | 5 | 0 | 6 | 1 | 0 | 0 | 0 | 16 |
| Egrandis | 196 | 0 | 0 | 7 | 0 | 0 | 0 | 0 | 1 | 1 | 57 | 0 | 370 | 4 | 178 | 327 | 1141 | 11 | 52 | 20 | 0 | 0 | 7 | 25 | 0 | 0 | 1 | 116 |
| Csinensis | 105 | 0 | 0 | 3 | 1 | 0 | 0 | 1 | 0 | 0 | 23 | 0 | 24 | 1 | 130 | 32 | 320 | 15 | 2 | 10 | 44 | 0 | 36 | 39 | 0 | 0 | 3 | 149 |
| Cclem | 198 | 0 | 0 | 43 | 2 | 0 | 0 | 0 | 1 | 0 | 38 | 0 | 118 | 4 | 261 | 81 | 746 | 36 | 11 | 48 | 80 | 0 | 452 | 183 | 0 | 0 | 4 | 814 |
| Ghirsutum | 157 | 0 | 0 | 8 | 35 | 0 | 0 | 0 | 3 | 0 | 133 | 0 | 1 | 0 | 158 | 19 | 514 | 10 | 378 | 15 | 18 | 1 | 184 | 110 | 0 | 0 | 14 | 730 |
| Cacao | 42 | 0 | 0 | 9 | 12 | 0 | 0 | 0 | 0 | 0 | 95 | 0 | 114 | 1 | 43 | 10 | 326 | 21 | 124 | 10 | 1 | 0 | 20 | 21 | 0 | 0 | 2 | 199 |
| papa | 53 | 0 | 0 | 0 | 0 | 0 | 0 | 0 | 0 | 0 | 27 | 0 | 0 | 0 | 2 | 17 | 99 | 8 | 37 | 0 | 6 | 0 | 0 | 0 | 0 | 0 | 1 | 52 |
| Thassleriana | 39 | 3 | 0 | 14 | 7 | 0 | 0 | 0 | 0 | 0 | 34 | 0 | 18 | 2 | 65 | 39 | 221 | 22 | 5 | 6 | 20 | 0 | 7 | 0 | 0 | 0 | 0 | 60 |
| Arabicum | 11 | 3 | 0 | 1 | 3 | 0 | 0 | 0 | 0 | 0 | 11 | 0 | 4 | 0 | 20 | 6 | 59 | 0 | 0 | 0 | 7 | 0 | 0 | 1 | 0 | 0 | 0 | 8 |
| Ath | 15 | 0 | 0 | 2 | 6 | 0 | 0 | 0 | 0 | 0 | 17 | 0 | 7 | 0 | 12 | 4 | 63 | 3 | 13 | 0 | 9 | 0 | 10 | 21 | 0 | 0 | 1 | 57 |
| Lalabamica | 6 | 0 | 0 | 0 | 0 | 0 | 0 | 1 | 0 | 0 | 13 | 0 | 1 | 0 | 5 | 2 | 28 | 2 | 1 | 0 | 16 | 0 | 0 | 0 | 0 | 0 | 0 | 19 |
| Eusalsugineum | 23 | 10 | 0 | 4 | 123 | 0 | 0 | 0 | 0 | 0 | 46 | 0 | 29 | 0 | 88 | 29 | 352 | 253 | 11 | 0 | 41 | 0 | 297 | 184 | 0 | 0 | 4 | 790 |
| Sparvula | 9 | 1 | 0 | 0 | 0 | 0 | 0 | 0 | 0 | 0 | 13 | 0 | 3 | 0 | 14 | 7 | 47 | 7 | 1 | 0 | 15 | 0 | 3 | 1 | 0 | 0 | 0 | 27 |
| Sirio | 24 | 11 | 0 | 0 | 2 | 0 | 0 | 0 | 0 | 0 | 13 | 0 | 9 | 0 | 9 | 18 | 86 | 12 | 0 | 0 | 8 | 0 | 0 | 0 | 0 | 0 | 0 | 20 |
| Brapa | 40 | 15 | 0 | 0 | 8 | 0 | 0 | 0 | 2 | 0 | 48 | 0 | 5 | 0 | 60 | 28 | 206 | 26 | 3 | 0 | 52 | 0 | 1 | 3 | 0 | 0 | 0 | 85 |
| Nnucifera | 276 | 2 | 2 | 11 | 0 | 0 | 0 | 0 | 1 | 0 | 15 | 0 | 0 | 0 | 26 | 119 | 452 | 44 | 0 | 1 | 89 | 0 | 9 | 276 | 0 | 0 | 3 | 422 |
| Acoerulea | 1 | 0 | 0 | 0 | 1 | 0 | 0 | 0 | 1 | 0 | 2 | 0 | 2 | 2 | 6 | 1 | 16 | 0 | 0 | 0 | 0 | 0 | 0 | 23 | 0 | 0 | 0 | 23 |
| Phoenix | 11 | 0 | 0 | 0 | 0 | 0 | 0 | 0 | 0 | 0 | 7 | 0 | 0 | 0 | 6 | 5 | 29 | 6 | 0 | 0 | 9 | 0 | 0 | 0 | 0 | 0 | 0 | 15 |
| oilpalm | 36 | 1 | 1 | 16 | 0 | 0 | 0 | 0 | 0 | 0 | 21 | 0 | 4 | 1 | 30 | 37 | 147 | 13 | 1 | 3 | 13 | 0 | 3 | 77 | 0 | 0 | 0 | 110 |
| banana | 5 | 0 | 0 | 5 | 0 | 0 | 0 | 0 | 0 | 0 | 13 | 0 | 30 | 0 | 76 | 11 | 140 | 16 | 5 | 22 | 109 | 0 | 0 | 14 | 0 | 0 | 0 | 166 |
| pineapple | 12 | 1 | 0 | 0 | 0 | 0 | 0 | 0 | 1 | 0 | 6 | 0 | 3 | 0 | 6 | 6 | 35 | 4 | 28 | 2 | 12 | 0 | 0 | 13 | 0 | 0 | 0 | 59 |
| Pedullis | 92 | 18 | 1 | 2 | 42 | 0 | 0 | 0 | 0 | 0 | 39 | 0 | 33 | 0 | 15 | 93 | 335 | 27 | 31 | 0 | 181 | 0 | 0 | 152 | 0 | 0 | 3 | 394 |
| Osativa | 59 | 19 | 0 | 3 | 5 | 0 | 0 | 0 | 0 | 0 | 75 | 0 | 63 | 0 | 175 | 13 | 412 | 34 | 280 | 0 | 93 | 0 | 6 | 749 | 0 | 0 | 10 | 1172 |
| Bdista | 47 | 18 | 0 | 30 | 16 | 0 | 0 | 0 | 0 | 0 | 55 | 0 | 16 | 0 | 27 | 28 | 237 | 29 | 22 | 0 | 37 | 0 | 0 | 239 | 0 | 0 | 0 | 327 |
| Taestivum | 159 | 67 | 0 | 263 | 10 | 0 | 0 | 0 | 0 | 0 | 50 | 0 | 144 | 0 | 45 | 255 | 993 | 203 | 363 | 0 | 129 | 1 | 148 | 1997 | 0 | 0 | 14 | 2855 |
| Pvirgatum | 35 | 22 | 0 | 7 | 4 | 0 | 0 | 0 | 3 | 0 | 16 | 0 | 21 | 0 | 50 | 18 | 176 | 8 | 17 | 0 | 57 | 1 | 22 | 137 | 0 | 0 | 14 | 256 |
| Etef | 38 | 11 | 0 | 0 | 2 | 0 | 0 | 0 | 0 | 0 | 4 | 0 | 0 | 0 | 0 | 13 | 68 | 1 | 0 | 0 | 34 | 0 | 1 | 1 | 0 | 0 | 0 | 37 |
| Sitalica | 69 | 38 | 0 | 391 | 1 | 0 | 0 | 0 | 0 | 0 | 45 | 0 | 16 | 0 | 91 | 142 | 793 | 163 | 93 | 0 | 51 | 8 | 1 | 704 | 0 | 0 | 147 | 1167 |
| Sbic | 18 | 22 | 0 | 0 | 10 | 0 | 0 | 0 | 0 | 0 | 56 | 0 | 79 | 0 | 22 | 12 | 219 | 44 | 143 | 0 | 45 | 1 | 133 | 549 | 0 | 0 | 3 | 918 |
| Zmays | 26 | 54 | 0 | 46 | 8 | 0 | 0 | 0 | 0 | 0 | 34 | 0 | 1737 | 0 | 34 | 18 | 1957 | 84 | 279 | 0 | 138 | 3 | 0 | 1964 | 0 | 0 | 119 | 2587 |
| Pequestris | 34 | 1 | 1 | 0 | 0 | 0 | 2 | 1 | 0 | 0 | 7 | 0 | 0 | 0 | 62 | 28 | 136 | 3 | 0 | 2 | 2 | 0 | 10 | 14 | 0 | 0 | 0 | 31 |
| Spolyrhiza | 17 | 0 | 0 | 0 | 0 | 0 | 0 | 0 | 0 | 0 | 24 | 0 | 0 | 0 | 5 | 8 | 54 | 5 | 0 | 10 | 13 | 0 | 8 | 0 | 0 | 0 | 0 | 36 |
| AmTr | 28 | 0 | 0 | 1 | 0 | 0 | 0 | 0 | 2 | 0 | 7 | 0 | 0 | 0 | 40 | 51 | 129 | 15 | 15 | 0 | 8 | 0 | 0 | 27 | 0 | 0 | 3 | 68 |
| Pabies | 5 | 0 | 0 | 0 | 0 | 0 | 1 | 126 | 0 | 0 | 44 | 0 | 15 | 0 | 26 | 133 | 350 | 143 | 57 | 0 | 86 | 0 | 29 | 79 | 0 | 0 | 8 | 402 |
| Selag | 0 | 0 | 0 | 0 | 0 | 0 | 17 | 0 | 0 | 0 | 0 | 0 | 0 | 0 | 0 | 23 | 40 | 0 | 41 | 228 | 4 | 0 | 32 | 12 | 0 | 252 | 96 | 665 |
| Ppatens | 0 | 0 | 0 | 0 | 1 | 44 | 0 | 0 | 0 | 1 | 0 | 0 | 0 | 0 | 1 | 13 | 60 | 0 | 0 | 3 | 0 | 0 | 264 | 18 | 121 | 0 | 70 | 476 |
| Creinhardtii | 0 | 0 | 0 | 0 | 0 | 0 | 0 | 0 | 0 | 0 | 0 | 4 | 0 | 0 | 0 | 0 | 4 | 0 | 0 | 0 | 0 | 0 | 0 | 0 | 0 | 0 | 9 | 9 |

Sup. Table 3B

| Specie | LARD | TRIM | TR_GAG |
|---|---|---|---|
| Bvulgar | 302 | 80 | 249 |
| Kiwifruit | 200 | 49 | 209 |
| Ecanadensis | 32 | 14 | 2 |
| Coffee | 257 | 46 | 383 |
| Smelongena | 105 | 40 | 38 |
| Slyco | 114 | 33 | 68 |
| Potato | 240 | 38 | 230 |
| Nicotiana | 295 | 18 | 209 |
| Cannuum | 386 | 39 | 208 |
| Mgut | 508 | 135 | 314 |
| Ugibba | 5 | 1 | 0 |
| Fexcelsior | 135 | 55 | 131 |
| Oeuropaea | 1010 | 392 | 240 |
| Vitis | 126 | 22 | 79 |
| Medicago | 133 | 11 | 90 |
| Cicer | 161 | 98 | 52 |
| Lotus | 91 | 28 | 123 |
| Ccajan | 208 | 46 | 217 |
| Phaseolus | 73 | 20 | 286 |
| Gmax | 151 | 65 | 542 |
| Araip | 718 | 50 | 517 |
| Csativa | 2311 | 72 | 675 |
| Malus | 456 | 49 | 37 |
| Fvesca | 116 | 10 | 30 |
| Pavium | 168 | 56 | 22 |
| PrunusP | 574 | 133 | 43 |
| watermelon | 16 | 10 | 4 |
| Cucumis | 7 | 4 | 1 |
| poplar | 62 | 17 | 7 |
| Rcommunis | 174 | 15 | 180 |
| Cassava | 81 | 37 | 27 |
| Hbrasiliensis | 235 | 39 | 143 |
| Lusitatissimum | 27 | 3 | 1 |
| Egrandis | 162 | 24 | 105 |
| Csinensis | 87 | 20 | 72 |
| Cclem | 96 | 36 | 145 |
| Ghirsutum | 110 | 13 | 371 |
| Cacao | 200 | 10 | 88 |
| papa | 27 | 10 | 13 |
| Thassleriana | 92 | 22 | 64 |
| Arabicum | 13 | 1 | 9 |
| Ath | 33 | 9 | 32 |
| Lalabamica | 23 | 1 | 3 |
| Eusalsugineum | 128 | 15 | 268 |
| Sparvula | 12 | 4 | 4 |
| Sirio | 25 | 10 | 6 |
| Brapa | 72 | 18 | 29 |
| Nnucifera | 131 | 13 | 90 |
| Acoerulea | 32 | 5 | 8 |
| Phoenix | 16 | 8 | 3 |
| oilpalm | 48 | 12 | 57 |
| banana | 112 | 15 | 98 |
| pineapple | 40 | 8 | 45 |
| Pedullis | 165 | 103 | 143 |
| Osativa | 466 | 40 | 262 |
| Bdista | 77 | 44 | 74 |
| Taestivum | 2541 | 119 | 2958 |
| Pvirgatum | 204 | 58 | 203 |
| Etef | 84 | 46 | 5 |
| Sitalica | 190 | 23 | 83 |
| Sbic | 249 | 25 | 342 |
| Zmays | 495 | 42 | 1465 |
| Pequestris | 71 | 36 | 58 |
| Spolyrhiza | 10 | 2 | 2 |
| AmTr | 89 | 52 | 25 |
| Pabies | 363 | 36 | 196 |
| Selag | 457 | 78 | 16 |
| Ppatens | 22 | 4 | 34 |
| Creinhardtii | 40 | 13 | 0 |

Sup Table 3C
