## Supplementary material for "High nucleotide similarity of three *Copia* lineage LTR retrotransposons among plant genomes": Sup Table 4

| Title | Authors | Year | Journal | Superfamily | Lineage | LTR-TE | Organisms | Methodology for HT | Transfer hypothesis | Total HT |
|---|---|---|---|---|---|---|---|---|---|---|
| Large distribution and | Dias *et al.* | 2015 | Plant | Copia | Unknown | *Copia25* | *Musa* (Musaceae) | High sequence | Mediated by | > 1 |
| Whole genome | Roulin *et al.* | 2009 | BMC | Unknown | Unknown | *Route66* | *Oryza* (Poaceae) and | High sequence | Mediated by | > 1 |
| EARE-1, a | Huang *et al.* | 2017 | Frontiers in | Copia | Angela/Tork | *EARE-1* | *Excoecaria* spp. | Phylogenetic | Mediated by viruses | 2 |
| Horizontal transfers of | Hou *et al.* | 2018 | Genome | Unknown | Unknown | *Mn.Zj.1 and* | *Morus notabilis* | Sequence | Mediated by bacteria, | 2 |
| Horizontal transfers of | Hou *et al.* | 2018 | Genome | Unknown | Unknown | *Md.Pb.1 and* | *Malus domestica* | Sequence | Mediated by bacteria, | 2 |
| Sample Sequence | Park, Christin & | 2021 | Molecular | Copia | Tork | Unknown | *Cymbopogon citratus* | High reverse | Mediated by insects | 2 |
| Sample Sequence | Park, Christin & | 2021 | Molecular | Gypsy | CRM | *CRM1* | *Echinochloa* | High reverse | Mediated by insects | 1 |
| Sample Sequence | Park, Christin & | 2021 | Molecular | Gypsy | CRM | *CRM1* | *Zuloagaea bulbosa* | High reverse | Mediated by insects | 1 |
| Sample Sequence | Park, Christin & | 2021 | Molecular | Gypsy | CRM | *wihov* | *Melinis ambigua* | High reverse | Mediated by insects | 1 |
| Sample Sequence | Park, Christin & | 2021 | Molecular | Gypsy | CRM | *wihov* | *Zuloagaea bulbosa* | High reverse | Mediated by insects | 1 |
| Sample Sequence | Park, Christin & | 2021 | Molecular | Gypsy | CRM | *wihov* | *Eriochloa meyeriana* | High reverse | Mediated by insects | 1 |
| Sample Sequence | Park, Christin & | 2021 | Molecular | Gypsy | CRM | *guhis* | *Echinochloa* | High reverse | Mediated by insects | 1 |
| Sample Sequence | Park, Christin & | 2021 | Molecular | Copia | Tork | *debeh* | *Echinochloa* | High reverse | Mediated by insects | 1 |
| Sample Sequence | Park, Christin & | 2021 | Molecular | Copia | Tork | *dounil* | *Echinochloa* spp. | High reverse | Mediated by insects | 2 |
| Sample Sequence | Park, Christin & | 2021 | Molecular | Copia | Retrofit | *lusi* | *Iseilema* | High reverse | Mediated by insects | 2 |
| Sample Sequence | Park, Christin & | 2021 | Molecular | Copia | Retrofit | *lusi* | *Cenchrus pilosus* | High reverse | Mediated by insects | 1 |
| Sample Sequence | Park, Christin & | 2021 | Molecular | Copia | Retrofit | *hera* | *Cenchrus setiger* | High reverse | Mediated by insects | 1 |
| Sample Sequence | Park, Christin & | 2021 | Molecular | Gypsy | CRM | 27 *CRM1* | *Echinochloa* spp. | Detection by | Mediated by insects | 27 |
| Sample Sequence | Park, Christin & | 2021 | Molecular | Gypsy | CRM | 9 *wihov* LTR- | *Echinochloa* spp. | Detection by | Mediated by insects | 9 |
| Sample Sequence | Park, Christin & | 2021 | Molecular | Gypsy | Unknown | 3 *pebi* LTR- | *Echinochloa* spp. | Detection by | Mediated by insects | 3 |
| Sample Sequence | Park, Christin & | 2021 | Molecular | Copia | Unknown | 21 *ibulaf* LTR- | *Echinochloa* spp. | Detection by | Mediated by insects | 21 |
| Sample Sequence | Park, Christin & | 2021 | Molecular | Copia | Unknown | 20 LTR-TEs | *Echinochloa* spp. | Detection by | Mediated by insects | 20 |
| Sample Sequence | Park, Christin & | 2021 | Molecular | Copia | Tork | 12 *dounil* | *Echinochloa* spp. | Detection by | Mediated by insects | 12 |
| Sample Sequence | Park, Christin & | 2021 | Molecular | Copia | Unknown | 5 *homy* LTR- | *Echinochloa* spp. | Detection by | Mediated by insects | 5 |
| Sample Sequence | Park, Christin & | 2021 | Molecular | Copia | Unknown | 3 *volo* LTR- | *Echinochloa* spp. | Detection by | Mediated by insects | 3 |
| Sample Sequence | Park, Christin & | 2021 | Molecular | Copia | Tork | 3 *debeh* LTR- | *Echinochloa* spp. | Detection by | Mediated by insects | 3 |
| Sample Sequence | Park, Christin & | 2021 | Molecular | Copia | Unknown | 2 *hani* LTR- | *Echinochloa* spp. | Detection by | Mediated by insects | 2 |
| Sample Sequence | Park, Christin & | 2021 | Molecular | Copia | Unknown | 2 *ijiret* LTR- | *Echinochloa* spp. | Detection by | Mediated by insects | 2 |
| Sample Sequence | Park, Christin & | 2021 | Molecular | Copia | Unknown | *fourf* | *Echinochloa* spp. | Detection by | Mediated by insects | 1 |
| Sample Sequence | Park, Christin & | 2021 | Molecular | Copia | Unknown | *tiwe* | *Echinochloa* spp. | Detection by | Mediated by insects | 1 |
| Horizontal transfer of | Park *et al.* | 2021 | International | Copia | Unknown | 30 LTR-TEs | *Vitis* spp. (Vitaceae) | High sequence | Mediated by fungi or | 30 |
| Horizontal transfer of | Park *et al.* | 2021 | International | Gypsy | Unknown | 4 LTR-TEs | *Vitis* spp. (Vitaceae) | High sequence | Mediated by fungi or | 4 |
| Evidence of multiple | Roulin *et al.* | 2008 | The Plant | Unknown | Unknown | *RIRE1* | Between *Oryza* | High sequence | (1) Mediated by | 7 |
| A new family of Ty1- | Cheng *et al.* | 2009 | Genetics | Copia | Unknown | *Rider* | *Arabidopsis* | High sequence | Mediated by | 2 |

Sup Table 4A

| Superfamily | Total HT Events | HT Events (%) |
|---|---|---|
| Copia | > 114 | 65,143 |
| Gypsy | 49 | 28,000 |
| Unknown | 12 | 6,857 |
| **Total** | **175** | **100,000** |

| Superfamily | Lineage | Total HT Events | HT Events (%) |
|---|---|---|---|
| Copia | *Angela/Tork* | 2 | 1,1429 |
| | *Retrofit* | 4 | 2,2857 |
| | *Tork* | 20 | 11,4286 |
| | Unknown | 88 | 50,2857 |
| Gypsy | *CRM* | 42 | 24,0000 |
| | Unknown | 7 | 4,0000 |
| Unknown | Unknown | 12 | 6,8571 |
| **Total** | | **175** | **100,0000** |

Sipe Table 4B
